# Melanoma antigens in pediatric medulloblastoma contribute to tumor heterogeneity and species-specificity of group 3 tumors

**DOI:** 10.1101/2024.05.14.594201

**Authors:** Rebecca R.J. Collins, Rebecca R. Florke Gee, Maria Camila Hoyos Sanchez, Sima Tozandehjani, Tara Bayat, Barbara Breznik, Anna K. Lee, Samuel T. Peters, Jon P. Connelly, Shondra M. Pruett-Miller, Martine F. Roussel, Dinesh Rakheja, Heather S. Tillman, Patrick Ryan Potts, Klementina Fon Tacer

**Author notes:** These authors contributed equally to this work (Rebecca R.J. Collins, Rebecca R. Florke Gee). Induced Proximity Platform, Amgen Research, Thousand Oaks, CA 91320, USA. Correspondence: Klementina Fon Tacer: School of Veterinary Medicine Texas Tech University, 7671 Evans Dr., Amarillo, TX 79106, ORCID: 0000-0003-1928-7745; Patrick Ryan Potts: Induced Proximity Platform, Amgen Research, Thousand Oaks, CA 91320.

## Abstract

**Background:** Medulloblastoma (MB) is the most malignant childhood brain cancer. Group 3 MB subtype accounts for about 25% of MB diagnoses and is associated with the most unfavorable outcomes. Herein, we report that more than half of group 3 MB tumors express melanoma antigens (MAGEs), which are potential prognostic and therapeutic markers. MAGEs are tumor antigens, expressed in several types of adult cancers and associated with poorer prognosis and therapy resistance; however, their expression in pediatric cancers is mostly unknown. The aim of this study was to determine whether *MAGEs* are activated in pediatric MB.

**Methods:** To determine *MAGE* frequency in pediatric MB, we obtained formalin-fixed paraffin-embedded tissue (FFPE) samples of 34 patients, collected between 2008 – 2015, from the Children’s Medical Center Dallas pathology archives and applied our validated reverse transcription quantitative PCR (RT-qPCR) assay to measure the relative expression of 23 *MAGE* cancer-testis antigen genes. To validate our data, we analyzed several published datasets from pediatric MB patients and patient-derived orthotopic xenografts, totaling 860 patients. We then examined how *MAGE* expression affects the growth and oncogenic potential of medulloblastoma cells by CRISPR-Cas9- and siRNA-mediated gene depletion.

**Results:** Our RT-qPCR analysis suggested that *MAGEs* were expressed in group 3/4 medulloblastoma. Further mining of bulk and single-cell RNA-sequencing datasets confirmed that 50-75% of group 3 tumors activate a subset of *MAGE* genes. Depletion of MAGEAs, B2, and Cs alter MB cell survival, viability, and clonogenic growth due to decreased proliferation and increased apoptosis.

**Conclusions:** These results indicate that targeting MAGEs in medulloblastoma may be a potential therapeutic option for group 3 medulloblastomas.

**Key Points:**

- Several Type I *MAGE* CTAs are expressed in >60% of group 3 MBs.
- Type I MAGEs affect MB cell proliferation and apoptosis.
- *MAGEs* are potential biomarkers and therapeutic targets for group 3 MBs.

**Importance of the Study:** This study is the first comprehensive analysis of all Type I *MAGE* CTAs (*MAGEA*, *-B*, and *-C* subfamily members) in pediatric MBs. Our results show that more than 60% of group 3 MBs express *MAGE* genes, which are required for the viability and growth of cells in which they are expressed. Collectively, these data provide novel insights into the antigen landscape of pediatric MBs. The activation of *MAGE* genes in group 3 MBs presents potential stratifying and therapeutic options.

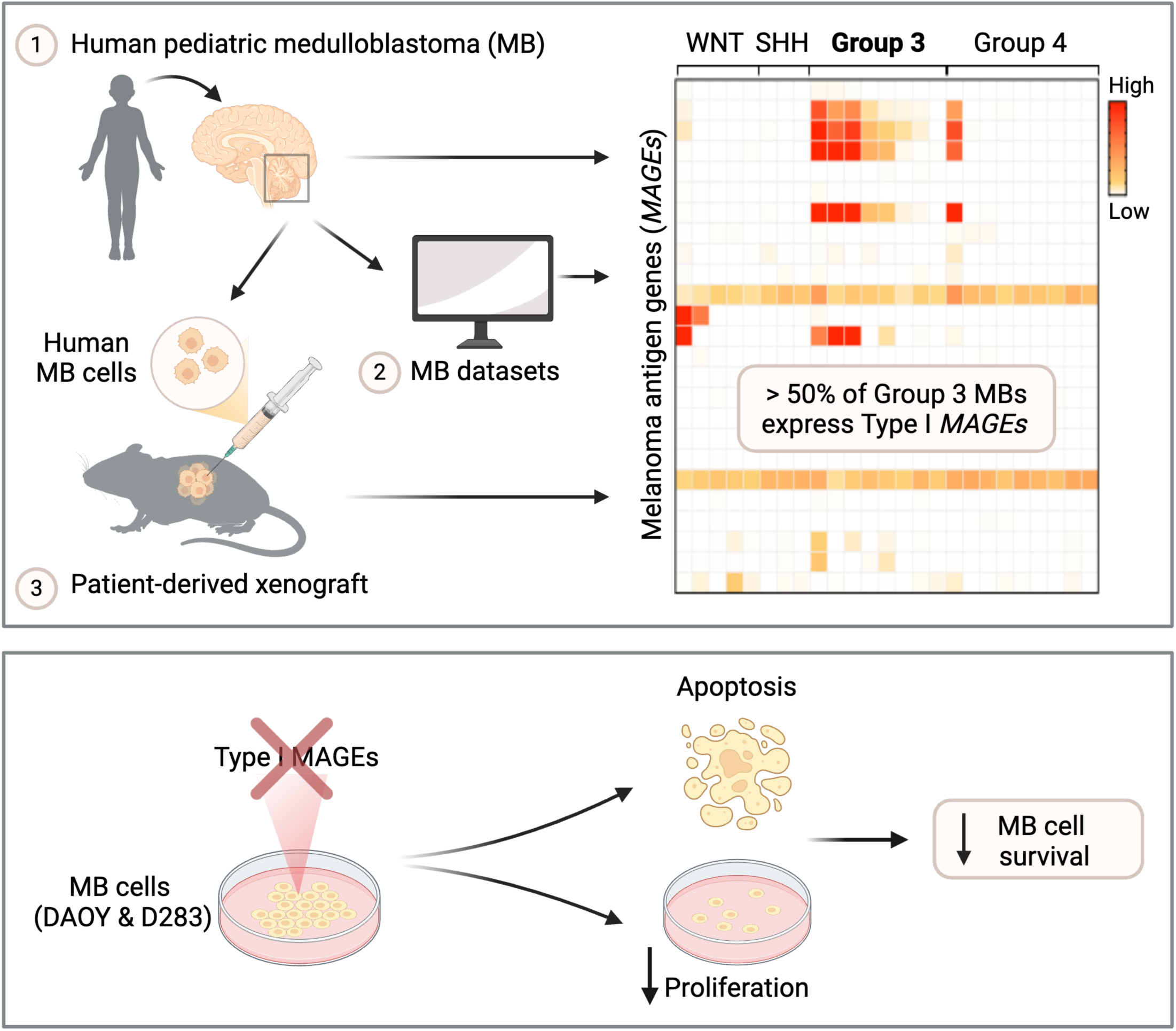

## Introduction

Medulloblastomas (MBs) are heterogeneous embryonal malignant tumors of the cerebellum that constitute one of the most common pediatric brain tumors with an incidence of six children per million under nine years of age (1-5). MBs are categorized into four molecular subgroups: Wingless (WNT), Sonic Hedgehog (SHH), group 3, and group 4 (5, 6). This categorization was first based on mutation profiling and expression arrays and, more recently, by DNA methylation (1, 7) and cellular origin (8, 9). Germline mutations in the WNT inhibitor *APC* or somatic mutations in *CTNNB1* are found in almost all WNT-MBs (10, 11). Etiology of SHH-MBs has been attributed to genetic changes in SHH signaling, including mutations in the SHH receptor *PTCH*, SHH inhibitor *SUFU*, or SHH transducer *SMO*, *GLI1/2* amplifications, and mutations in *MYCN* and *TP53* (8, 10, 12). Group 3 and group 4 MBs (Group 3/4), which account for about 60% of MB diagnoses, are the least understood with respect to disease biology and developmental origins, as they exhibit complex and sometimes overlapping mutational spectrums, DNA methylation profiles, and expression characteristics (5-8, 13). Moreover, group 3 tumors are associated with the worst prognosis of all the subgroups and are frequently metastatic at presentation (6), calling for novel approaches for effective diagnosis, prognosis, and treatment of patients with these tumors.

Surgery, radiation, and chemotherapy have improved MB patients’ prognosis in the last decades, but approximately 30% of patients remain incurable, and survivors suffer severe long-term side effects from these therapies. Distinct molecular signatures of the 4 subgroups enables a personalized therapeutic approach that is already benefiting patients (2). However, among the four subgroups, group 3 MBs exhibit the highest incidence of high-risk characteristics and therapy resistance (7). Over the last decade, immunotherapy has rapidly changed the therapeutic landscape and prognosis for many adult cancers and is also in development for pediatric solid tumors (14). Immunotherapies for MB have begun to undergo clinical testing (ClinicalTrials.gov: NCT03173950, NCT01326104); thus, identification of selective targets, biomarkers of responsiveness, and strategies for overcoming resistance in these tumors will be of utmost importance.

Although several brain tumor-specific targets have been identified, only a limited number of studies have investigated antigen expression and validity in pediatric populations (15). To prevent toxicities, the best targets for immunotherapy are tumor-associated antigens that are exclusively expressed in tumor cells with minimal expression in normal tissues (16). Tumor-associated antigens are commonly neo-antigens generated by tumor cells because of genomic mutations (17). Given that the somatic mutation burden increases with patient age, childhood cancers have a lower mutation burden and mutation-derived neo-antigen levels (18). Intriguingly, tumors also commonly activate the expression of genes normally restricted to male germ cells, referred to as cancer-testis antigens (CTAs), as expression outside of their naturally immune-privileged site in the testis can activate an immune response (19). Melanoma-associated antigens (MAGEs) were the first CTAs discovered (20-22). The *MAGE* family encompasses approximately 40 conserved genes divided into two major subgroups: Type I and Type II (20). While Type II *MAGEs* are ubiquitously expressed and implicated in neurodevelopment, Type I *MAGEs* are CTAs with normal expression in the testis but aberrantly expressed in various cancers (20, 21, 23, 24). Furthermore, Type I MAGEs predict poor patient prognosis and are remarkable candidates for immunotherapy targets (20-25). MAGEs are heavily investigated in immunotherapy of adult cancers (25), but very little is known about their expression and role in pediatric tumors (26, 27).

In this study, we performed the first comprehensive analysis of all Type I *MAGE* CTAs (*MAGEA*, *-B*, and *-C* subfamily members) in pediatric MBs and found that several are expressed in more than 60% of group 3 MBs and are required for the viability and growth of cells in which they are expressed. Collectively, these data provide novel insights into the antigen landscape of pediatric MBs and show that more than half of group 3 tumors activate *MAGE* genes, presenting potential stratifying and therapeutic options.

## Materials and Methods

### Materials

Refer to Extended Materials and Methods.

### Medulloblastoma patient samples

With appropriate institutional review board approval, we searched for brain tumor pathology cases diagnosed as medulloblastoma with a histologic subtype during the years 2008-2015 that also had FFPE tissue archived at Children’s Medical Center Dallas. Paraffin blocks and glass slides were examined to confirm the diagnosis and ensure adequate tissue availability. 34 de-identified cases were selected, and all except MT23 had sufficient material for complete analysis. Scrolls 10 μm thick were obtained from at least one block of tissue for each case. Genetic subtyping of the medulloblastoma cases was done using paired expression of DKK1 and WIF1 (>0.01 value of both genes corresponds to WNT subtype) or SFRP and HHIP (>0.01 value of both genes corresponds to SHH subtype); all other cases were classified as Group 3/4 subtype. Genetic subtyping was confirmed by immunohistochemistry staining profile in 8 cases (Table S1). No cases had an immunoprofile that refuted the genetic subtype (one case was unable to be classified by immunostains).

### Ethics approval and consent to participate

The present study was performed in accordance with the guidelines proposed in the Declaration of Helsinki and with institutional approval. FFPE tissue collection and phenotypic and tissue analyses were conducted under UT Southwestern Institutional Review Board approval IRB STU 102015-047. MT17 cells were cultured with patient/family consent under Pediatric Biospecimen Repository protocol IRB STU 082010-115. MFR research was conducted under IRB NBTPO1-UT-XPD. All animal studies were approved by the Animal Care and Use Committee and performed in accordance with best practices outlined by the NIH Office of Laboratory Animal Welfare.

### Cell culture

DAOY (cat. no. HTB-186) and D283 (cat. no. HTB-185) cells were purchased from American Type Culture Collection (ATCC) and grown in DMEM supplemented with 10% FBS, 2 mM L-glutamine, and 1x antibiotic-antimycotic (100 units/mL penicillin, 100 mg/mL streptomycin, and 0.25 mg/mL Amphotericin B). At the time of patient surgery, MT17 tumor cells were obtained from the Pediatric Biospecimen Repository; these cells were collected as residual tissue from the patient’s surgical resection specimen, cultured in minimum essential media (MEM) supplemented with 10% FBS and 10% BM-Condimed (Roche) at the affiliated hospital, and then cryogenically stored. We subsequently cultured MT17 cells in DMEM supplemented with 10% FBS, 2mM L-glutamine, 1x antibiotic-antimycotic, and 10% BM-Condimed. All cells were incubated in a humidified atmosphere at 37°C with 5% CO_2_. Cell counts were obtained with a hemocytometer or with a Countess II Automated Cell Counter using trypan blue staining.

### RNA isolation and reverse transcription quantitative PCR (RT-qPCR)

RNA from FFPE tissues was isolated using the Qiagen RNeasy FFPE Kit according to the manufacturer’s protocol. RNA from cell cultures, mouse tumors, and PDOX samples was isolated using the TRIzol reagent (Invitrogen) according to the manufacturer’s directions. RNA was treated with DNase I (Roche) in 4.5 mM MgCl_2_ to eliminate genomic DNA contamination, and cDNA was prepared from 4 μg of DNase-treated RNA using the Applied Biosystems High-Capacity cDNA Reverse Transcription (Applied Biosystems) kit in 100 μL final volume. Following cDNA synthesis, RNase-free water was added to increase the sample volume to 300 μL. Gene expression levels were measured by RT-qPCR in triplicate wells of a 384-well reaction plate with 22 ng of cDNA/well on an Applied Biosystems 7900HT with SYBR Green chemistry. As previously published (23), specific primers were used for each Type I *MAGE* (150 nM concentration; Table S3). Relative gene expression was calculated by normalizing against 18S values and calibrated across different plates. Extended Materials and Methods provides additional details.

### MAGE KO Library

Up to five sgRNAs for each *MAGE* and *MAGE*-related gene were designed and assessed for off-target potential *in silico* based off homology to other sites in the genome. sgRNAs were cloned into the lentiGuide-Puro vector (Addgene #52963). The finished library consisted of 494 sgRNAs targeting 155 human and mouse genes, along with non-targeting negative controls making up 2.5% of the library (Table S2). Validation to check sgRNA presence and representation was performed using calc_auc_v1.1.py (https://github.com/mhegde/) and count_spacers.py (28). Viral particles were produced by the St. Jude Vector Laboratory.

### CRISPR dropout screen

The generation of DAOY-Cas9 stable expressing cells is described in Extended Materials and Methods. For the screen, 40,000 DAOY-Cas9 stable expressing cells were seeded in 10 wells of 6-well plates. The next day, cells were transduced with the MAGE KO library (MOI < 0.5) in the presence of 8 µg/mL (final concentration) polybrene (Sigma-Aldrich). 48 h after viral transduction, 500,000 cells were collected as the Day 0 sample for NGS analysis, and the remaining cells were selected with 0.5 µg/mL puromycin. Cells were split when about 80-90% confluent, and at least 500,000 cells were collected at this time by washing with PBS, pelleting cells, and freezing cells at -80 °C. At the end of 30 days, genomic DNA was extracted from the samples with the DNEasy Blood & Tissue Kit (Qiagen) following manufacturer’s protocol. St. Jude Center for Advanced Genome Editing (CAGE) PCR amplified the integrated sgRNAs as described in the Broad GPP protocol (https://portals.broadinstitute.org/gpp/public/resources/protocols). The St. Jude Hartwell Center Genome Sequencing Facility provided NGS sequencing using single end 100 cycle kit on a NovaSeq 6000 (Illumina). NGS data were analyzed using MAGeCK-VISPR/0.5.7 (29). Results are reported as log_2_ fold change for Day 30 sgRNA reads using the Day 0 reads as the baseline, and datapoints represent triplicate samples for all sgRNAs targeting a specific gene. The abundance of each sgRNA over time was determined by setting the Day 0 abundance as 100%.

### siRNA transfection and viability assays

For knockdown experiments, cells were reverse transfected with 33.3 nM siRNA using Lipofectamine RNAiMAX (Invitrogen) according to the manufacturer’s instructions. Cells were then plated in 6-well plates, and the media was changed 24 h after transfection. Cells were collected 48-72 h after transfection for qPCR or western blotting. Control siRNAs for LonRF (oligo # 3012482021-000160, -000170) and UBB (oligo # 3030837381-000300, -000310) were purchased from Sigma. Sequences of all other siRNAs (Sigma-Aldrich) are included in Table S3.

Cells were seeded 3,000-5,000 cells/well in 96-well plates and grown for 72-96 h following reverse transfection. AlamarBlue (Bio-Rad) or CellTiter-Glo Luminescent Cell Viability Assay (Promega) was used to measure cell viability, along with an automated plate reader that measured fluorescence or luminescence, respectively. Results are reported as a percentage normalized to the averaged siRevL1 control. Experiments were repeated up to three times on separate days.

### BrdU immunofluorescence assay

4,000 DAOY cells/well were plated in 8-well chambered glass slides (Nunc Lab-Tek II) and reverse transfected with siRNA. After 48 h, 10 µM (final concentration) of BrdU (BD Pharmingen BrdU FlowKit 559619) was incubated with cells for 4 h. Then, cells were washed with ice-cold PBS, fixed with methanol for 10 min at -20°C, and washed with room temperature (RT) PBS. Cells were incubated in 2 M hydrochloric acid for 20 min at RT to dissociate dsDNA, washed with PBS, and permeabilized with blocking solution [PBS containing 0.2% (v/v) Triton X-100 and 3% (w/v) bovine serum albumin (BSA)] for 20 min at 4°C. Anti-BrdU rat antibody conjugated to FITC (ab74545; 1:250 or 1:500) was incubated with cells at RT for 1 h in the dark. After washing with PBS containing 0.2% Triton X-100, nuclei were stained with DAPI. Stained cells were then mounted with Aqua-Poly/Mount (Polysciences) and imaged at 20x with a Leica AF6000 microscope and appropriate filters. For each slide, an untreated well with and without BrdU was used to establish background fluorescence. Images were analyzed with ImageJ and BrdU-positive cells determined by counting at least 100 cells for each condition.

### Clonogenic assays

DAOY cells were reverse transfected in 6-well plates and allowed to grow overnight. Cells were then trypsinized, diluted, mixed with an equal volume of warm agar (0.35% final), and 1 x 10^4^ cells/well placed in 6-well plates over a 0.5% agar base. After cooling for 20 min, 1 mL culture media was added to the top of each well. After 24 h, individual cells were counted within 10 separate fields using an inverted microscope with a 20x objective, to ensure equal cell densities between wells (data not shown). Plates were incubated for 4 weeks with media changed 1-2 times a week. DAOY colonies were imaged with a 4x objective and all colonies >50 micrometers were counted in 9 separate fields. The soft agar assay with D283 cells and the colony formation assay is described in Extended Materials and Methods.

### Preparation of cell lysates and western blotting

Cell lysates were prepared as previously described (30), and the total protein concentration quantified with the Micro BCA Protein Assay Kit (Thermo Scientific). Lysates were prepared in SDS sample buffer, resolved on SDS-PAGE gels, and transferred to nitrocellulose membranes. Membranes were blocked with 5% BSA in TBST [25 mM Tris pH 8.0, 2.7 mM KCl, 137 mM NaCl, 0.05% (v/v) Tween-20] and incubated with primary antibodies. The following antibodies were used: anti-MAGE-B2 (30), anti-MAGE-A2 (31), anti-Cleaved PARP (Asp214) (19F4) (Cell Signaling Technology, #9546), anti-TRIM28 (Abcam, ab22553), anti-Cas9 (Abcam, ab191468), and anti-β-actin (Abcam, ab6276). After three washes with TBST, membranes were incubated with secondary antibodies, washed an additional three times, and detected via chemiluminescence using ECL detection reagent (GE Healthcare, RPN2209).

### MB microarray and single-cell RNA-sequencing (scRNA-seq) datasets

GlioVis data portal (32) was used to analyze and visualize Cavalli et al. (33) data of 763 MB patient samples (GSE85218). We obtained the scRNA-seq datasets of primary MB patient samples and patient-derived xenografts from GSE119926 (8) and GSE155446 (34). Average expression of *MAGE* genes was calculated for all the cells of each patient/xenograft to generate heatmaps. Additional details are provided in Extended Materials and Methods.

### Plotting and statistical analyses

Except Figures S1, S2B-F, and S3, plots were made in GraphPad Prism 10. Results are expressed as the mean ± standard deviation from at least two independent experiments with individual data points indicated. Significance was assessed with one-way or two-way ANOVA followed by Dunnett’s multiple comparisons test for all samples compared to the control [*P* ≤ 0.05 (∗), *P* ≤ 0.01 (∗∗), *P ≤* 0.001 (∗∗∗), *P* ≥ 0.05 (non-significant, ns)].

## Results

### Type I MAGE CTAs are expressed mainly in group 3 of pediatric medulloblastoma

To determine the frequency of *MAGE* expression in pediatric MB, we measured expression levels of Type I *MAGEs* in patient tumors (Figure 1A). We obtained FFPE tissue samples from 34 patients, collected between 2008 and 2015, from the Children’s Medical Center Dallas pathology archives and performed RT-qPCR. This cohort of MB samples contained one unknown, one WNT, 11 SHH, and 21 group 3/4 molecular subtypes (Table S1). To confirm the subgroups, we measured WNT inhibitors [WNT Inhibitor Factor 1 (WIF1) and Dickkopf 1 (DKK1)] and SHH-target genes (SFRP1 and HHIP) known to be produced by WNT- and SHH-MBs, respectively (35, 36). Although type I *MAGE*s are not expressed in normal brain tissues (Figure S4F)(23), 66% of the group 3/4-MBs expressed at least one *MAGE* gene, with many tumors expressing multiple *MAGEs* (Figure 1B). In contrast, the WNT- and SHH-MBs were positive for only one or two *MAGEs* (except for MT29 which expressed three). Our analysis suggested disease-specific expression of Type I *MAGEs* in group 3/4 MBs, in line with their common expression in more aggressive adult tumors (20).

**Figure 1.**
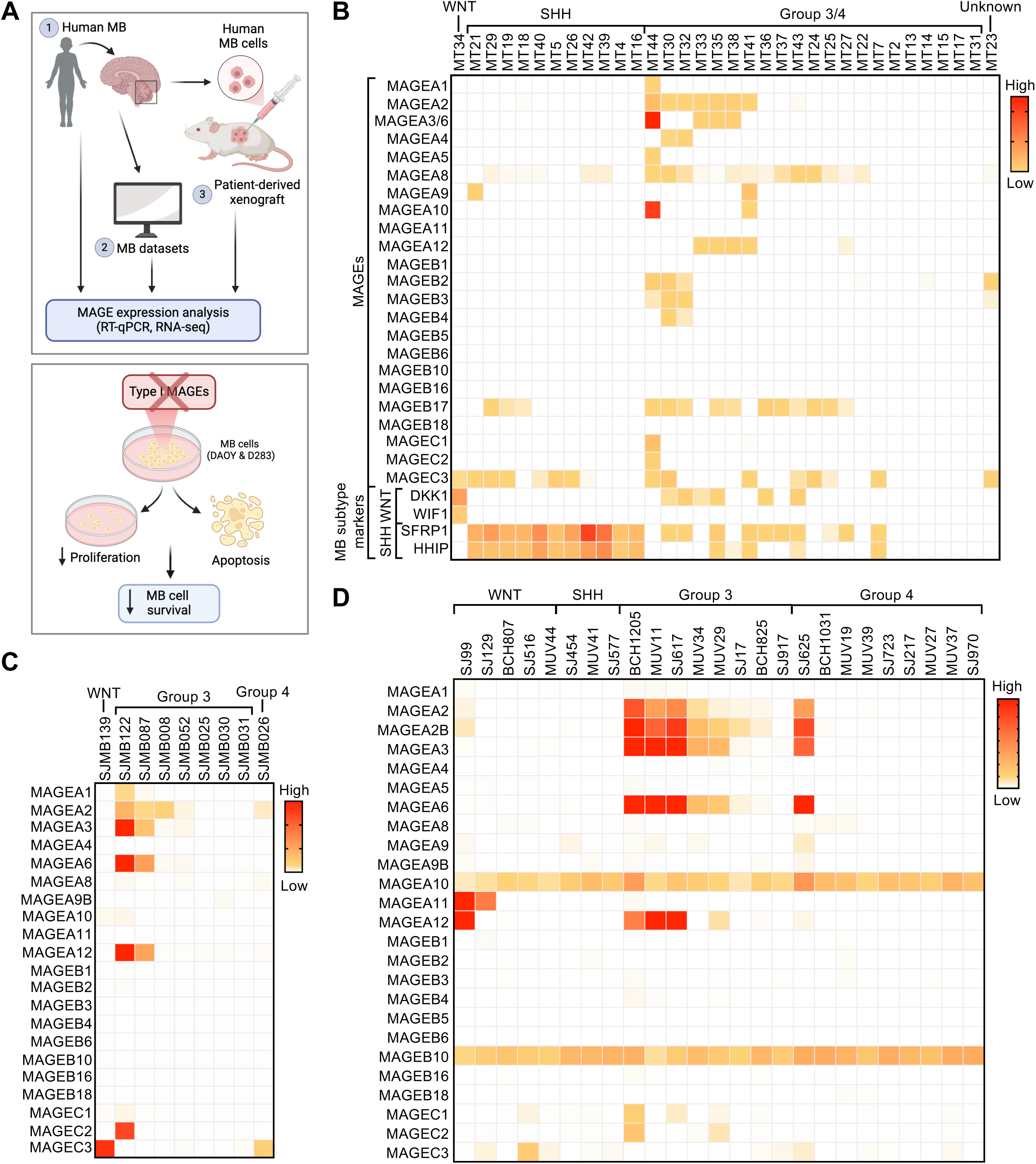
Group 3 medulloblastomas express multiple Type I *MAGEs*. **(A)** Overview of data presented herein. Heatmaps show the expression of *MAGEs* in pediatric medulloblastomas with indicated subtypes, as determined by **(B)** qPCR (Pediatric Biospecimen Repository at Children’s Medical Center Dallas), **(C)** RNA-seq (Pediatric Cancer Genome Project, data downloaded 2.4.2020), or **(D)** scRNA-seq [GSE119926 dataset (8)].

To validate our findings, we analyzed publicly available and published datasets for *MAGE* expression (Figure 1A). We first examined the GlioVis database (32) for *MAGE* mRNA levels in 763 pediatric MBs analyzed by microarray (33) and found that *MAGEA1, -A3, -A11, -B2,* and -*C* subfamily members were detected (data not shown). *MAGEA3* expression was highly enriched in group 3, and its expression was linked to worse survival (Figures S1A and S1B). The expression of other Type I *MAGEs*, like *MAGEB2*, did not show any specific pattern (Figure S1C), except *MAGEC3* enrichment in the WNT subgroup (Figure S1D). We then specifically looked for RNA sequencing (RNA-seq) data, as microarray probes are often not specific enough to distinguish between several genes in the *MAGEA, -B,* and -*C* subfamilies that exhibit high levels of similarity (20). Our analyses of bulk RNA-seq data from the PeCan database (St. Jude Children’s Research Hospital; N=9; Figure 1C) and scRNA-seq data published by Hovestadt et al. (8) (N=25; Figure 1D) and Riemondy et al. (34) (N=29; Figure S2A) revealed a similar trend in *MAGE* expression in MB tumors (Figure 1B). By combining the single-cell expression for each tumor, we found that more than half of all the tumors analyzed by scRNA-seq were positive for > 3 *MAGEs*, with group 3 MBs exhibiting the highest expression levels of multiple Type I *MAGEs* (Figure 1D). These data support our initial finding that *MAGE* CTAs are expressed in a significant portion of patients with group 3 MBs and may represent a stratifying marker for this heterogeneous group of patients.

### Patient-derived orthotopic xenograft (PDOX) models recapitulate tumor MAGE expression signature

We then analyzed the expression of *MAGE* genes in patient-derived orthotopic xenograft (PDOX) models of childhood brain tumors, generated from pediatric brain tumor patients treated at St. Jude Children’s Research Hospital (Figure 1A) (37). RT-qPCR and RNA-seq analyses revealed that these PDOX models at least partially recapitulated the patterns of *MAGE* expression seen in patients. In particular, the majority of group 3 PDOX models expressed multiple Type I *MAGEs* (Figures 2A and 2B). Similarly, our analysis of a published PDOX scRNA-seq dataset (8) showed that 66% of the samples express at least one *MAGE*, and group 3 PDOX models are positive for multiple *MAGEs* (Figure 2C). These findings demonstrate that several Type I *MAGEs* are also expressed in PDOX models derived from group 3 MBs.

**Figure 2.**
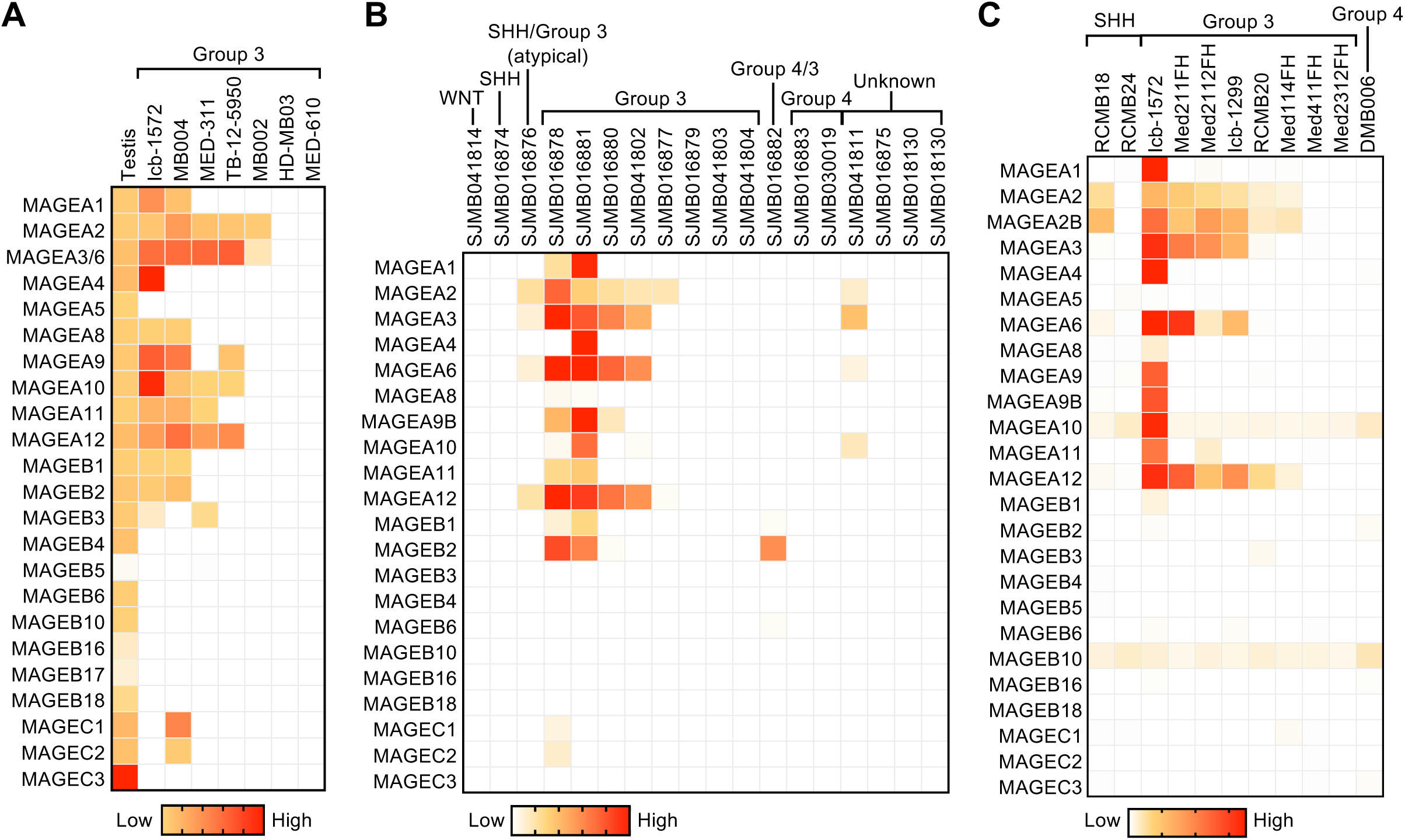
Patient-derived orthotopic xenograft (PDOX) models from group 3 medulloblastomas express multiple *MAGEs*. Heatmaps show the expression of *MAGEs* in PDOX models, as determined by **(A)** RT-qPCR [generated in Dr. Martine Roussel lab (37)] **(B)** RNA-seq [generated and analyzed in Dr. Martine Roussel lab (37)], or **(C)** scRNA-seq [GSE119926 dataset (8)].

### MAGEs are expressed in distinct subsets of cells in MAGE-positive group 3 MB tumors

To determine the heterogeneity of MAGE expression in each tumor, we analyzed *MAGE* expression in MB tumors on a single-cell level. In the dataset published by Riemondy et al. (34), most malignant cells were grouped according to different individuals, while myeloid cells, lymphocytes, and oligodendrocytes were not separated between patients (Figure S2B), alluding to interindividual differences in MB tumors. In line with the ubiquitous expression pattern reported for most Type II *MAGEs* (Figure S4F) (23), these genes are expressed in all cell types without showing a specific pattern, as shown for *MAGED2* expression distribution between neoplastic and non-neoplastic cells (Figure S2C). However, expression of Type I *MAGEs* (i.e., *MAGEA3, -A10*, and -*A12*) was restricted to malignant cells from group 3 MBs and to specific patients (Figures S2D-F). For example, patient 1130 expressed *MAGEA3, -A6*, and *-A12*, with some cells from patients 1355 and 1028 also expressing *MAGEA3* and *-A6* (Figures S2D, S2F, and S3C). On the other hand, the expression of *MAGEA10* was restricted to patient 1167 (Figure S2E). All these samples had representation of different subpopulations of neoplastic cells (mitotic, undifferentiated progenitor, and neuronally differentiated); however, the expression of *MAGEs* was not restricted to specific subpopulations.

From the scRNA-seq dataset published by Hovestadt et al. (8), we determined the percentage of cells from the *MAGE*-positive group 3 MBs expressing each Type I *MAGE* (Table S4) and the number of Type I *MAGEs* expressed (Table S5). We found that *MAGEA2/2B, -A3, -A6, -A10, -A12*, and -*B10* were expressed in at least 20% of cells in three or more group 3 MB samples (Table S4). In all but one group 3 MB sample, the majority of cells expressed one or two *MAGEs* (Table S5).

Further, we wanted to know whether these *MAGEs* are activated in the same cells or distinct cell groups. Our clustering visualization of all *MAGE*-expressing cells from all the patients suggested that different *MAGE* genes are often expressed in different subsets of cells (Figures S3A and S3B) (8, 34). This pattern is also apparent when zoomed in on individual patients (Figure S3C) (34) and as also seen for *MAGEA3* and *-A10* in Figures S2D and S2E. Given that the functions of individual MAGE proteins may be discrete, this mosaic *MAGE* expression may contribute to tumor heterogeneity.

### MAGE expression contributes to the species-specificity signature of group 3 MBs

To understand tumor initiation and identify tumor-specific therapeutic targets, previous studies have focused on identifying the cellular origins of childhood MB (8, 38, 39). Comparison of human cerebellar tumors with the developing murine cerebellum indicated that WNT, SHH, and group 3 tumors consisted of subgroup-specific undifferentiated and differentiated neuronal-like malignant populations, whereas group 4 tumors were exclusively comprised of differentiated neuronal-like neoplastic cells (8, 39). The tumor cells expressing *MAGEs* likely comprise undifferentiated progenitor-like cells with high MYC activity (subpopulation program B), as Hovestadt et al. (8) reported that more than 88% of the cells from the group 3 MBs, in which multiple *MAGEs* were expressed (Figures 1D, 2C, and S3A), were annotated as program B.

The lack of high-confidence correlations between murine cerebellar populations and group 3 MBs (8, 39) suggests a cell of origin for group 3 MB that is absent in the developing murine cerebellum. Although the mouse cerebellum shares many features of lamination, circuitry, neuronal morphology, and foliation with humans, the human cerebellum has 750-fold greater surface area, increased neuronal numbers, altered neuronal subtype ratios, and increased folial complexity, suggesting species-specific neuronal progenitors (40). Indeed, Smith et al. (9) recently reported that group 3 MBs are closely aligned with human fetal rhombic lip progenitors. Intriguingly, we could not detect expression of any Type I *Mages* in tumors from diverse MB mouse models [i.e., cMyc overexpression (*Myc* OE), *Myc* overexpression with co-deletion of *p53* and *p18* (*Trp53*^−/−^, *Cdn2c*^−/−^, MYCN), co-deletion of *Ptch* and *p53* (*Ptch1*^+/-^, *Trp53*^-/-^)] (data not shown) (41). These data suggest that Type I *MAGE* genes have species-specific oncogenic potential and expression regulation, which is in line with the recent evolution and expansion of Type I *MAGE* genes, oftentimes in a species-specific manner (20, 23).

### Disrupted expression of MAGEs decreases the viability of medulloblastoma cells

To determine whether MAGEs contribute to MB cell viability, as reported in other cancer cell types (24, 42-44), we evaluated *MAGE* expression profiles in MB cell lines (Figure 3A), many of which express multiple Type I *MAGEs* like the MB tumor and PDOX samples (Figures 1 and 2). DAOY and D283 MB cell lines were selected for experiments after confirming their patterns of *MAGE* expression (Figures 3A, S4A, and S4G). First, we generated DAOY-Cas9 stable expressing cells and confirmed Cas9 activity (Figures S4B and S4C). Then, we conducted a CRISPR dropout screen in DAOY-Cas9 cells transduced with a lentiviral pooled library containing non-targeting sgRNAs (negative controls) and approximately five sgRNAs targeting each *MAGE* gene (Figure 3B). Many of the sgRNAs targeting the ubiquitously expressed Type II *MAGEs* (Figures S4F and S4G), particularly *NSMCE3* (*MAGEG1*) and *MAGEL2*, were depleted by day 30 (Figures S4D and S4E), which supports previous reports on their importance for cell viability (20, 45, 46). In addition, many of the sgRNAs targeting the Type I *MAGE* CTAs (Figure S4F) yielded negative CRISPR scores, including *MAGEA3/6* and -*B2* (Figures 3C and 3D). Besides specific sgRNAs that only targeted one *MAGE* gene, the pooled library also contained promiscuous sgRNAs that targeted multiple *MAGEs* with similar sequences. Interestingly, the 14 sgRNAs targeting *MAGEA* subfamily members were depleted over time, with many showing reductions similar to or lower than that of the specific *MAGEA3/6* sgRNAs (Figure 3E).

**Figure 3.**
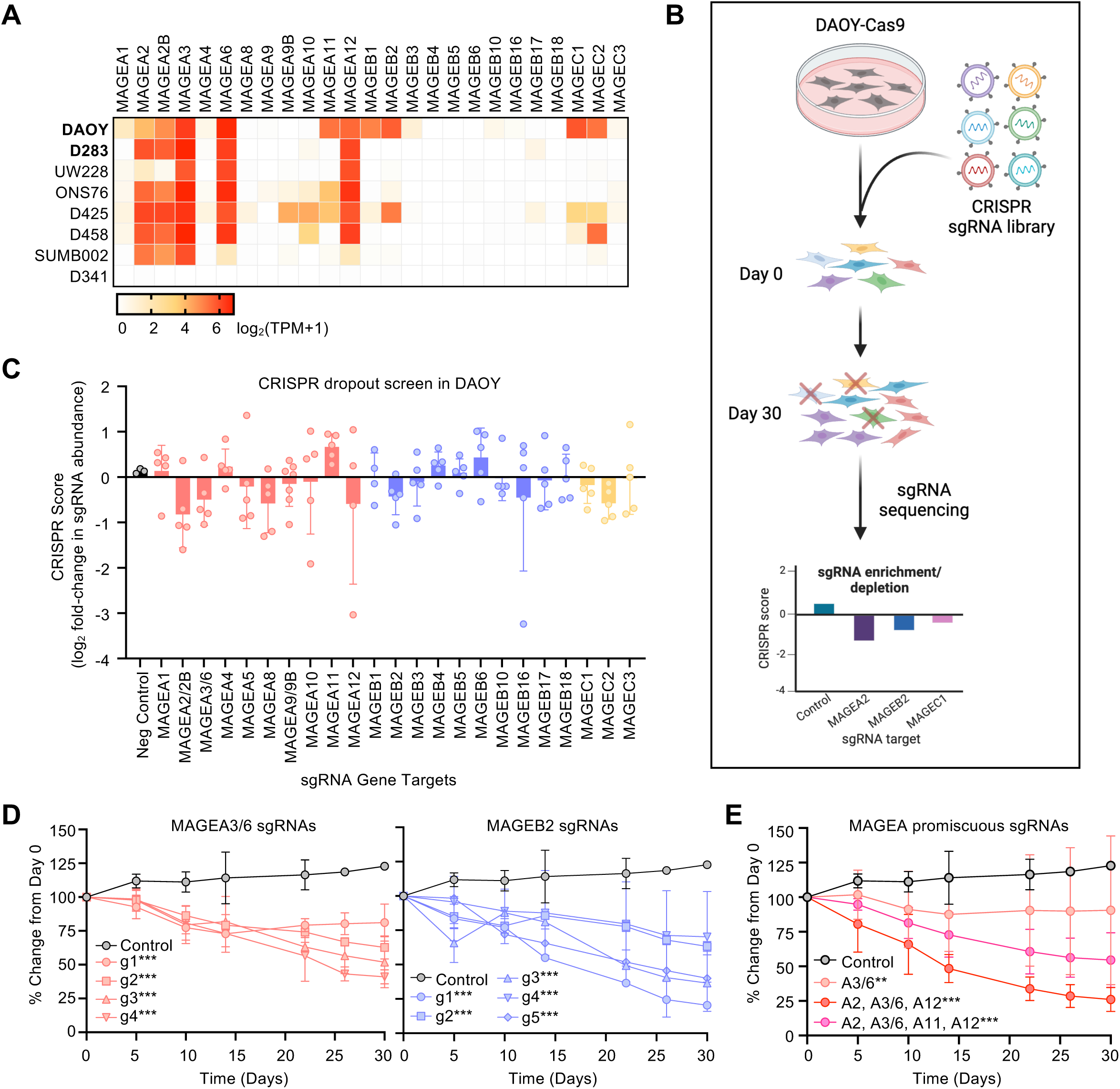
Depleted MAGE expression decreases the viability of DAOY medulloblastoma cells. **(A)** Expression of Type I *MAGEs* in medulloblastoma cell lines (data downloaded from DepMap Public 23Q2 on 10.11.2023). Values are inferred from RNA-seq data using the RSEM tool and are reported after log_2_ transformation, using a pseudo-count of 1; log_2_(TPM+1). **(B)** Experimental schematic of CRISPR dropout screen performed in DAOY-Cas9 stable expressing cells. **(C)** CRISPR score (log_2_ fold change from day 0) was calculated for the abundance of all sgRNAs on day 30. Data points show the average CRISPR score of each sgRNA targeting a particular Type I MAGE (n = 3). **(D)** Percent change is shown over time for each sgRNA targeting *MAGEA3/6* (*left*) or *MAGEB2* (*right*). Data points show the average percent change for each sgRNA from triplicate samples. **(E)** Promiscuous sgRNAs targeting multiple MAGEAs were depleted over time. Data points show the average percent change from day 0 for all sgRNAs targeting the same group of multiple *MAGEA* genes (n = 3). Significance was assessed with two-way ANOVA followed by Dunnett’s multiple comparisons test for samples compared to the non-targeting control sgRNAs [*P* ≤ 0.01 (∗∗), *P* ≤ 0.001 (∗∗∗)].

As another approach to determine which MAGE proteins contribute to cell viability, we transiently knocked down Type I *MAGEs* in DAOY and D283 MB cells using siRNAs. The siRNA-mediated knockdown of Type I *MAGEs* decreased MB cell viability (Figures 4A, 4B, and S5A). We confirmed the reduced expression of siRNA targets by RT-qPCR (Figure S4H), with siPanMAGEA2 and -A4 affecting the predicted *MAGEAs* (Figures S4I). Interestingly, DAOY cell viability was reduced the most upon knockdown of *MAGEB2* (Figures 4B and S5A). Cell viability was minimally affected by knocking down *MAGEs* in MT17 MB cells that did not express Type I *MAGEs*, suggesting that the effect on cell viability in DAOY cells was dependent on *MAGE* expression (Figures 1A, 4C, and 4D).

**Figure 4.**
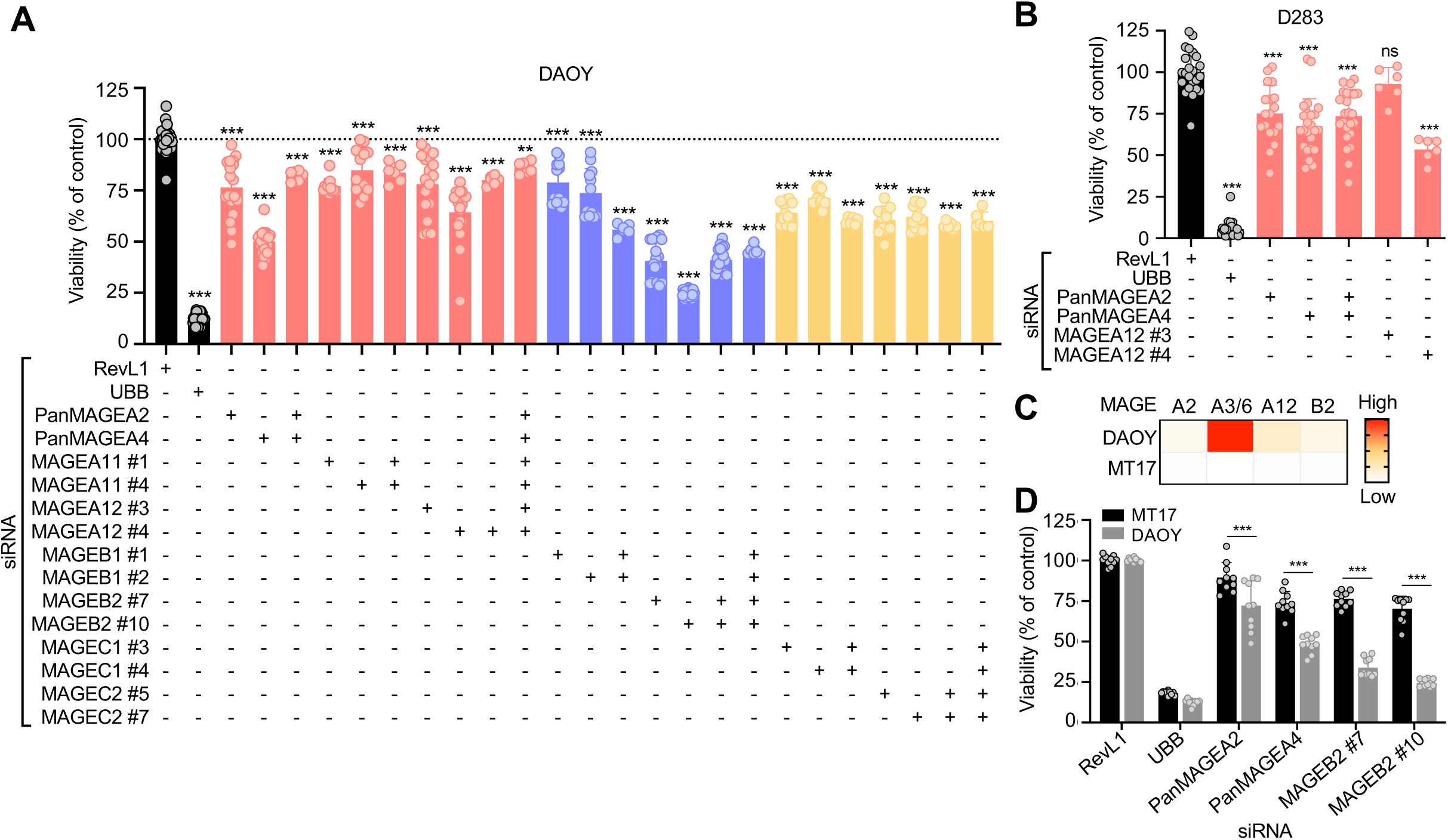
Knockdown of MAGEs decreases the viability of medulloblastoma cells. **(A)** DAOY or **(B)** D283 medulloblastoma cells were transfected with the indicated siRNA(s), and AlamarBlue or CellTiter-Glo viability assay, respectively, was performed after 3 days. The viability percentage was calculated by normalizing to the siRevL1 control. Graphs show normalized data points for at least 4 replicates from 2-6 experiments. Significance was assessed with one-way ANOVA followed by Dunnett’s multiple comparisons test for samples compared to the siRevL1 control [*P* ≤ 0.05 (∗), *P* ≤ 0.01 (∗∗), *P* ≤ 0.001 (∗∗∗), *P* > 0.05 (non-significant, ns)]. **(C)** Heatmap shows expression of selected Type I *MAGEs*, as determined by RT-qPCR, in DAOY cells compared to MT17 cells. **(D)** Compared to MT17 cells, the siRNA-mediated knockdown of the indicated MAGEs significantly reduces DAOY cell viability, as determined by two-way ANOVA. Relevant DAOY data was extracted from **A** for comparison with MT17, as the experiments could not be done at the same time.

### MAGEs contribute to medulloblastoma cell proliferation and growth

The decreased cell viability observed upon depletion of MAGEA and MAGEB2 proteins (Figures 3C-3E, 4A, 4B, and S5A) suggested that MAGEs may play a role in cell proliferation. Indeed, siRNA-mediated knockdown of MAGEAs and MAGEB2 moderately decreased the percentage of BrdU-positive cells, particularly in cells transfected with siPanMAGEA4, which knocked down all MAGEAs (Figures 5A and 5B). Next, we evaluated whether Type I MAGEs participate in anchorage-independent growth as a marker of malignancy. The number of colonies formed in soft agar by DAOY cells transfected with MAGEA or -B2 siRNAs was significantly reduced compared to control cells (Figures 5C, 5D, S5B, and S5C), indicating that these MAGEs contribute to tumorigenicity. The siRNA-mediated depletion of MAGEAs and -B2 in D283 cells also led to decreased colony formation (Figures S5D and S5E). These reductions in cell proliferation and colony growth upon MAGE knockdown may be due to increased cell death by apoptosis or necrosis, at least for MAGEB2 knockdown, which led to the production of cleaved PARP (Figure 5E) and a higher percentage of propidium iodide (PI)-positive cells (Figures S5F and S5G). Together, these data suggest that expression of Type I *MAGEs* in MB cancer cells leads to dependence on their function(s), and MAGE depletion decreases cell viability or even leads to cell death.

**Figure 5.**
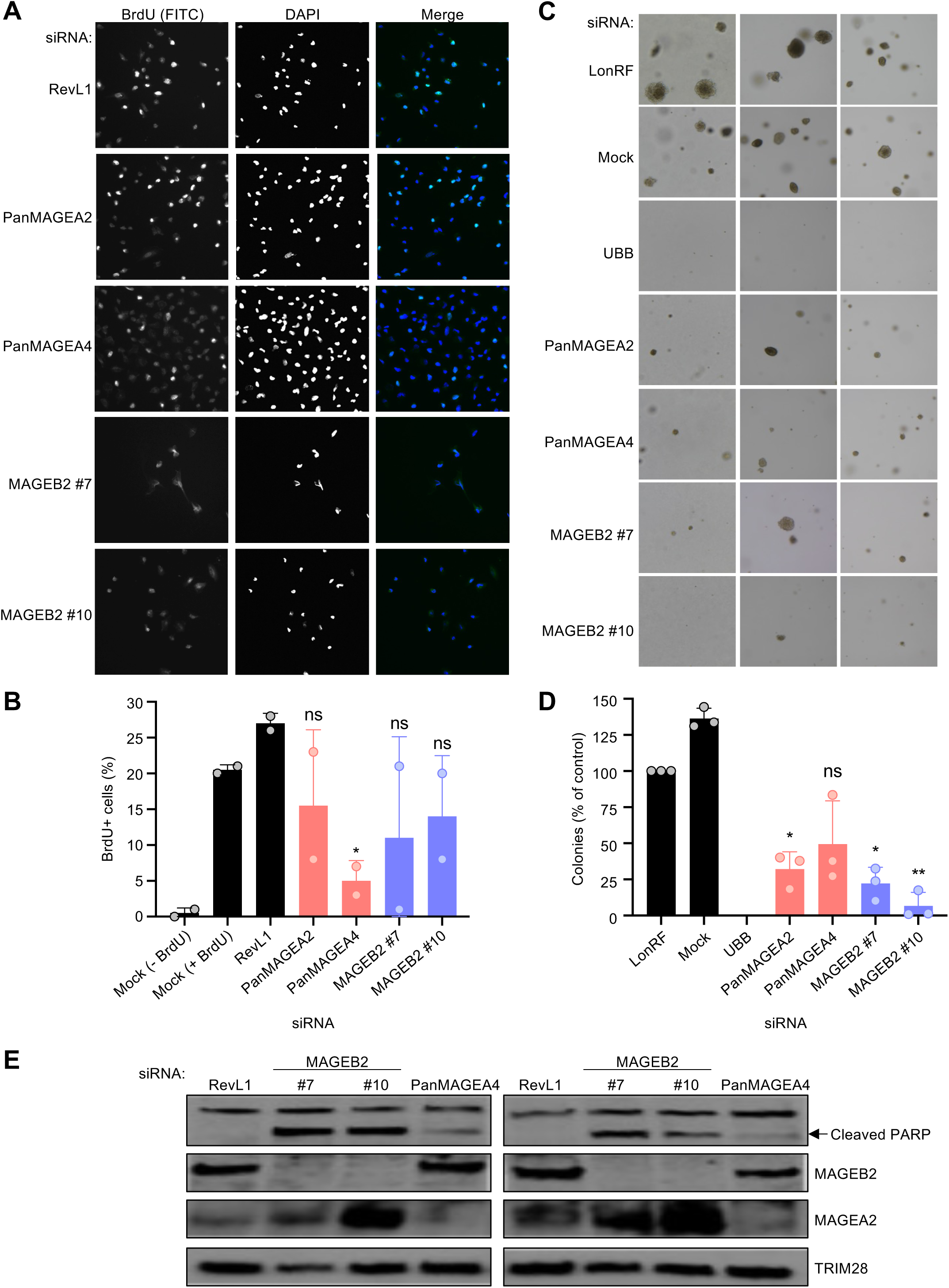
Knockdown of MAGEs decreases colony formation and cell proliferation and increases apoptosis. **(A)** DAOY cells were transfected with the indicated siRNA and BrdU immunofluorescence assay was performed 48 h later. Representative images of stained cells are shown. **(B)** Graph shows the percentage of BrdU-positive cells, relative to the siRevL1 control, determined by counting at least 100 cells for each condition (n = 2). **(C)** Transfected DAOY cells were plated for a soft agar assay, and colonies were imaged after 4 weeks. Representative images for each siRNA are shown. **(D)** The number of colonies for each condition was normalized to the siLonRF control to calculate the percentage (n = 3). **(E)** DAOY cells were transfected with indicated siRNA and collected 48 h after transfection for western blotting. siMAGEB2s and, to a lesser extent, siPanMAGEA4 increased cleavage of PARP, indicative of increased apoptosis. Significance was assessed with one-way ANOVA followed by Dunnett’s multiple comparisons test for samples compared to the siRevL1 or siLonRF control [*P* ≤ 0.05 (∗), *P* ≤ 0.01 (∗∗), *P* > 0.05 (non-significant, ns)].

## Discussion

The tumor antigen landscape of pediatric MB remains largely unexplored. This report is the first comprehensive analysis of the expression of all Type I *MAGEs* in pediatric MBs and their association with MB subgroup, as earlier studies did not report MB subtype or were classified differently than the current consensus (10). Consistent with previous analyses of individual *MAGE-A* and *-C* family members (26, 27, 47, 48), we found that at least one Type I *MAGE* is expressed in more than half of pediatric MBs (Figure 1). *MAGE* expression was particularly evident in group 3 MBs, which is considered the most aggressive MB subgroup, with a 5-year overall survival of less than 60% (49). Although group 3 MBs exhibit some notable genomic features, such as specific aneuploidies and amplification of *MYC, MYCN*, or *OTX2*, the unresolved molecular heterogeneity of these tumors contributes to the low therapeutic success reported in patients (11, 49, 50).

We discovered that more than 60% of group 3 MBs express at least one, but often several, *MAGE* genes (Figures 1, 2, and S1-S3). Different *MAGE* genes are often expressed in distinct subsets of cells (Figures S2, S3, and Tables S4-S5), suggesting they contribute to the intra- and inter-tumor heterogeneity of MBs. However, further studies are needed to identify the epigenetic and transcriptional mechanisms responsible for activating aberrant *MAGE* expression in MBs. The co-expression of multiple Type I *MAGEs* in neoplastic cells may impart oncogenic functions, such as increased survival and proliferation, due to synergistic molecular effects or these proteins acting together (51). Accordingly, *MAGE* expression in various cancers is associated with chemoresistance and a worse prognosis (52-57). In the testis, subsets of *MAGE* genes are expressed in distinct populations of male germ cells during differentiation, starting from spermatogonial stem cells through haploid spermatids, suggesting autocrine and paracrine functions (23). Importantly, we previously reported that *MAGE* genes protect the germline against diverse stressors (*e.g.*, nutritional stress, chemotherapy, and heat), indicating that their protective functions may have evolved to protect male fertility but are hijacked by cancer cells to increase their growth and therapy resistance (23, 30). Therefore, *MAGE* expression signature may contribute to better stratification of and therapy selection for group 3 patients.

Targeting these aberrantly expressed *MAGEs* or the pathways they regulate may represent potential therapeutic opportunities, including immunotherapy. Their restricted tissue expression (23) may lead to a lower probability of side effects and toxicity, critically important in the pediatric population that is still undergoing brain development (49). The decreased viability, proliferation, and colony formation we observed upon depletion of Type I *MAGEs* (Figures 3-5) suggests that these genes play a role in the proliferation and viability of MB cells and supports targeting *MAGEs* as a potential mode of MB cancer therapy. However, further investigation into the molecular functions of these *MAGEs* in MBs is needed. The expression of *MAGEs* in group 3 is especially attractive, as current therapeutic options are limited and have a high risk of neurotoxicity (10, 33, 58). Targeting multiple MAGEs concurrently may further allow for a synergistic effect of therapy while reducing the side effects.

Additionally, we found that the *MAGE* expression profile in PDOX models mirrored the tumors (Figure 2). Given that the group 3 cell of origin is human-specific and not found in mice (9), these PDOX models represent an important experimental model for studying MAGEs in group 3 MBs. In contrast to adult cancers, which are frequently attributable to genomic alterations due to repeated environmental exposures, childhood malignancies are more often the consequence of failed developmental processes (59). We previously found that several Type I *MAGEs* are expressed more broadly during embryonic development, particularly in the brain, indicating they may be involved in developmental processes and, when derailed, contribute to oncogenic transformation (23). Prototypic group 3 MBs are predominantly comprised of undifferentiated progenitor-like cells with high MYC activity (8), suggesting that the undifferentiated transcriptional program of these cells may be connected to their aberrant expression of multiple *MAGEs*. Furthermore, the undetectable expression of *Mages* in tumors from murine genetic MB models further corroborates the species-specificity of this subtype and calls for enhanced development of patient-derived models, including PDOXs and organoids, to substitute for laboratory rodents.

In conclusion, we report that Type I *MAGEs* are expressed in more than half of pediatric MBs, particularly in group 3 tumors. Further investigation into their tumor-intrinsic functions and impact on the immune microenvironment is needed, as MAGEs represent an exciting opportunity to address the biggest unmet needs for group 3 MB patients: rigorous preclinical biomarkers and therapeutic targets. The depletion of *MAGEs* led to decreased viability, proliferation, and colony formation in MB cell lines, suggesting that MAGE-targeted therapy could enhance chemosensitivity in *MAGE*-positive MBs. Further preclinical research is needed to better understand potential therapeutic vulnerabilities of *MAGE*-positive group 3 MBs and how these can be preclinically tested using more accurate models.

## Additional Information

### Authorship

RRJC, PRP, and KFT conceived and designed the research. RRJC, RRFG, AKL, MCHS, ST, TB, and BB conducted experiments and analyzed data. RRJC, RRFG, MCHS, TB, ST, and KFT wrote the manuscript. MFR, STP, JPC, and SMP-M developed reagents, and DR provided resources for this study. All authors have approved the final manuscript.

### Data availability

Data generated in this work are available upon reasonable request. Publicly available data accession numbers are reported above in the Materials and Method section.

## Acknowledgments

We thank members of the Fon Tacer and Potts laboratories for their advice and critical discussions. The graphical abstract and Figures 1A and 3B were created with BioRender (BioRender.com).

