## Supplemental material for "Melanoma antigens in pediatric medulloblastoma contribute to tumor heterogeneity and species-specificity of group 3 tumors"

**Supplementary Material**

### Extended Materials and Methods

#### Materials

Lipofectamine RNAiMAX transfection reagent, L-glutamine and antibiotic-antimycotic stock solutions, blasticidin, puromycin, phosphate-buffered saline (PBS), Opti-MEM reduced serum medium, and Dulbecco's Modified Eagle's Medium (DMEM) were all purchased from Thermo Fisher Scientific (Atlanta, GA, USA). Fetal bovine serum (FBS) was purchased from HyClone (Logan, UT, USA). AlamarBlue was purchased from Bio-Rad (Hercules, CA, USA), and CellTiter-Glo Luminescent Cell Viability Assay was purchased from Promega (Madison, WI, USA). Polybrene and crystal violet were purchased from Sigma-Aldrich (St. Louis, MO, USA). All tissue culture supplemental materials, such as tissue culture flasks and disposable pipettes, were purchased from Corning (Corning, NY, USA).

#### CRISPR dropout screen

To generate DAOY-Cas9 stable expressing cells, 40,000 DAOY cells were seeded in a 6-well plate. The next day, cells were transduced with Cas9 lentivirus (MOI <1) in the presence of 8 µg/mL (final concentration) polybrene. Forty-eight hours after transfection, cells were selected with 2.5 µg/mL blasticidin. After selecting cells for 2–4 weeks, stable expression of Cas9 protein was confirmed by western blot (Figure S4B). Cells were continuously cultured in the presence of 2.5 µg/mL blasticidin after selection. Cas9 activity was evaluated by targeted next-generation sequencing (NGS), as previously reported (29), three days after transfecting parental DAOY and DAOY-Cas9 cells with 300 ng eGFP plasmid (Lonza) and 700 ng sgRNA (Synthego) targeting *F9* via nucleofection (Lonza 4D-Nucleofector, X unit) with P3 solution and the EN-158 program.

#### RT-qPCR analysis

Cycling conditions were as follows: 50°C for 2 minutes, 95°C for 10 minutes, and 40 cycles of 95°C for 15 seconds followed by 60°C for 60 seconds. Control reactions were performed without RT (−RT) to ensure that amplifications were from RNA and not contaminating genomic DNA. *MAGEA2/A2B*, *-A3/A6*, and *-A9/A9B* genes are too similar for specific primers and thus primers that detect both genes were used (23). For primer validation and efficiency determination, standard cDNA generated from mouse/human testis RNA was used as described (23). Briefly, PCR efficiencies were calculated from the slope of the resulting standard curves. FFPE brain tissue from an autopsy case was used as a normal control. RT-qPCR data were analyzed by ABI instrument software SDS2.3. Baseline values of amplification plots were set automatically, and threshold values were kept constant to obtain normalized cycle time and linear regression data. Relative gene expression was calculated by dividing the averaged, efficiency-corrected values for mRNA expression by that for 18S ribosomal RNA, multiplying the quotient by $1 \times 10^5$ for scaling purposes, and calibrating across different plates. Expression levels were defined as absent if the cycle time value was more than 34. Results with SD values greater than the expression value were rejected.

#### Clonogenic assays

The soft agar assay with D283 cells was conducted as described for DAOY cells, except plates were incubated for 5 weeks. D283 colonies were stained with 0.5 mL/well of crystal violet for >1 h at room temperature, washed with water, and dried before counting all stained colonies visible by unaided eye.

For the colony formation assay, DAOY cells were transfected with siRNA in 6-well plates. After 24 h, cells were trypsinized, and 500 cells/well were plated in 6-well plates in 2 mL media. Plates were incubated for 12 days with media changes every 2-3 days. Then, colonies were fixed and stained with crystal violet solution (0.05% w/v crystal violet, 1% formaldehyde, 1% methanol in PBS) for 20 min, washed with water, and dried. Stained colonies were imaged using Azure Systems 600 and quantified using ImageJ software. Results are reported as a percentage of colonies relative to the siLonRF control.

**Apoptosis by flow cytometry**

Cells are reverse transfected (50nM siRNA) in 6-well cell culture plates and allowed to grow for 48 h. Cells are then carefully collected and stained with Annexin V (Molecular Probes Inc. Dead Cell Apoptosis Kit with Annexin V AlexaFluor 488; AV) and propidium iodide (PI). Cells were analyzed by flow cytometry by gating for viable cells on the forward and side scatter. Then, single-stained untreated controls were used to separate PI-/AV-, PI-/AV+, PI+/AV+, PI+/AV- cells into quadrants. Approximately 20,000 live cells were counted. Results are reported as fold change relative to the siRevL1 control for each quadrant.

**Medulloblastoma microarray and single-cell RNA-sequencing (scRNA-seq) datasets**

All patients and MB subtypes from datasets were included. From GSE119926 (8), the primary patient samples dataset included 25 samples (23 diagnostic and 2 recurrences) and 7745 malignant cells in total. The patient-derived xenografts dataset included 11 samples and 946 cells in total. The dataset from GSE155446 (32) included 28 samples and 39946 cells in total, and visualization of *MAGE* gene expression via T-SNE plots in these samples was obtained from UCSC Cell Brower (https://www.pneuroonccellatlas.org/project/mb/).

Additionally, from the single-cell datasets (8, 32), cells with at least one Type I *MAGE* gene expressed (expression level > 0) were filtered in an Excel (Microsoft Office) table. Then, a clustered heatmap was generated based on *MAGE* gene expression in these cells using the heatmap.2 function from the gplots package in R (R Core Team (2022) R: A Language and Environment for Statistical Computing. R Foundation for Statistical Computing, Vienna, https://www.R-project.org). Similarity was calculated through the distfun parameter for rows (distfunRow) and columns (distfunCol) using dist() function. The distance was calculated by default with Euclidean distance (method = "euclidean") which calculates the straight-line distance between two points in Euclidean space, or for a pair of vectors, it computes the square root of the sum of squared differences between corresponding elements of the vectors. Using the same filter and approach, a heatmap was made for individual samples (Figure S3).

### Supplemental Figure Legends

**Supplemental Figure 1.** *MAGEA3* is expressed in group 3 MB tumors **(A)** and is associated with poorer prognosis **(B)**. Expression of *MAGEA3* **(A)**, *MAGEB2* **(C)**, and *MAGEC3* **(D)** were analyzed and plotted by GlioVis (30) data portal for visualization and analysis of brain tumor expression datasets of 763 MB patient samples [GSE85218 (31)].

**Supplemental Figure 2. (A)** Heatmap shows the average expression of Type I *MAGEs* in MBs, as determined by scRNA-seq [GSE155446 (32)]. **(B)** Unaligned UMAP projection of single-cell expression data from 28 MB patient samples. Expression of **(C)** *MAGED2*, **(D)** *MAGEA3*, **(E)** *MAGEA10*, and **(F)** *MAGEA12* in cells from patients with MB. Cells from Group 3-MBs are circled in red. Figures B-F were obtained from UCSC Cell Browser (https://www.pneuroonccellatlas.org/project/mb/) on 10.17.2023.

**Supplemental Figure 3. (A-B)** Expression of *MAGE* genes in *MAGE*-positive cells from scRNA-seq datasets. **(C)** Expression pattern of *MAGE* genes in *MAGE*-positive cells from individual pediatric MBs. Data in **A** is from GSE119926 (8), and data in **B** and **C** is from GSE155446 (32).

**Supplemental Figure 4. (A)** Expression of Type I *MAGEs* in DAOY and D283 cells was confirmed by qPCR. **(B)** Western blot shows that Cas9 is expressed in DAOY-Cas9 stable expressing cells. **(C)** Cas9 activity was evaluated by transfecting parental DAOY and DAOY-Cas9 stable expressing cells with sgRNA targeting *F9* and collecting for targeted NGS after 3 days. Increased percentage of total indels in *F9* gene in DAOY-Cas9 cells indicates functional Cas9 activity. **(D)** CRISPR score ($log_2$ fold change from day 0) was calculated for the abundance of all sgRNAs on day 30. Data points show the average CRISPR score of each sgRNA targeting a particular type II *MAGE* (n = 3). Significance was assessed with one-way ANOVA followed by Dunnett's multiple comparisons test for samples compared to the negative control sgRNAs [$P \leq 0.001$ (∗∗∗)]. **(E)** Data points show the average percent change over time for each sgRNA targeting *NSMCE3* (*left*) or *MAGEL2* (*right*) (n = 3). **(F)** Heatmap shows restricted expression pattern of Type I *MAGEs* in testis and ubiquitous expression pattern of Type II *MAGEs* across most tissues [RNA-seq data downloaded from the Genotype-Tissue Expression (GTeX) portal on 1.16.2024]. **(G)** Expression of Type II *MAGEs* in medulloblastoma cell lines (data downloaded from DepMap Public 23Q2 on 10.11.2023). Values are inferred from RNA-seq data using the RSEM tool and are reported after $log_2$ transformation, using a pseudo-count of 1; $log_2$(TPM+1). **(H)** Relative expression for the indicated *MAGEs* was determined by RT-qPCR two days after transfecting DAOY cells with specified siRNAs. **(I)** Heatmap shows the predicted MAGEs affected by siPanMAGEA2 and -A4 based on sequence (*left*), and graph shows expression of indicated *MAGEAs*, determined by RT-qPCR, after siRNA transfection in DAOY cells (*right*).

**Supplemental Figure 5. (A)** DAOY medulloblastoma cells were transfected with the indicated siRNA, and AlamarBlue viability assay performed after 3 days. Viability percentage was calculated by normalizing to the siRevL1 control (n = 6). **(B)** Transfected DAOY cells were plated for a colony formation assay, and colonies imaged after 12 days. **(C)** The number of DAOY colonies for each condition was normalized to the siRevL1 control to calculate the percentage (n = 3). **(D)** Transfected D283 cells were plated for a soft agar assay, and colonies were imaged after 5 weeks. **(E)** The number of D283 colonies for each condition was normalized to the siLonRF control to calculate the percentage (n = 3). **(F)** DAOY cells were transfected with the indicated siRNA and after 48 h, stained with Annexin V (AV) and propidium iodide (PI). Cells were analyzed by flow cytometry by gating for viable cells on the forward and side scatter and then separated into quadrants using single-stained untreated controls. Approximately 20,000 live cells were counted.

**(G)** Flow cytometry results are reported as fold change relative to the siRevL1 control for each quadrant. Significance was assessed with one-way ANOVA followed by Dunnett's multiple comparisons test for samples compared to the siLonRF control [$P \leq 0.05$ (∗), $P \leq 0.01$ (∗∗), $P > 0.05$ (non-significant, ns)].

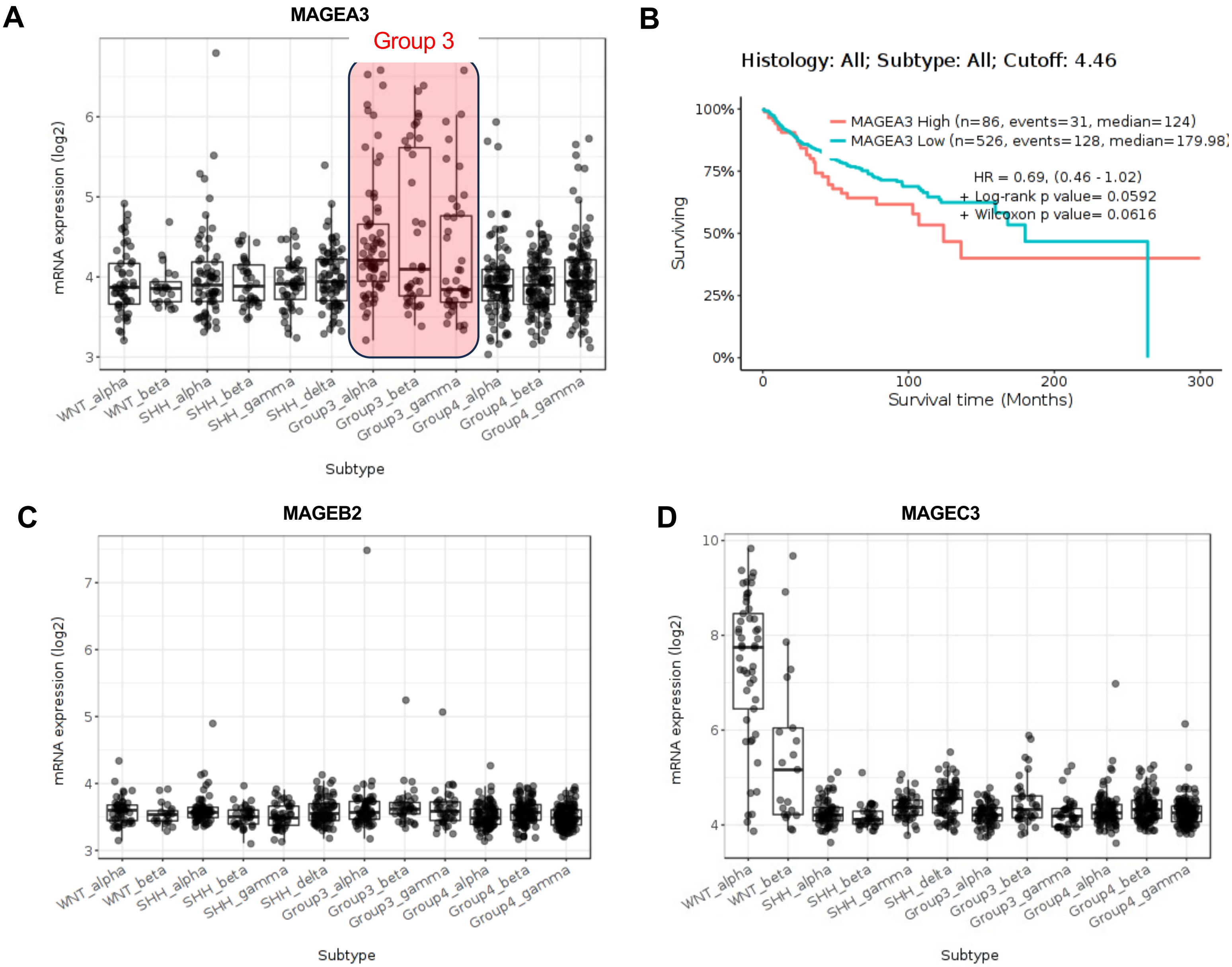


**Supp. Figure 1**

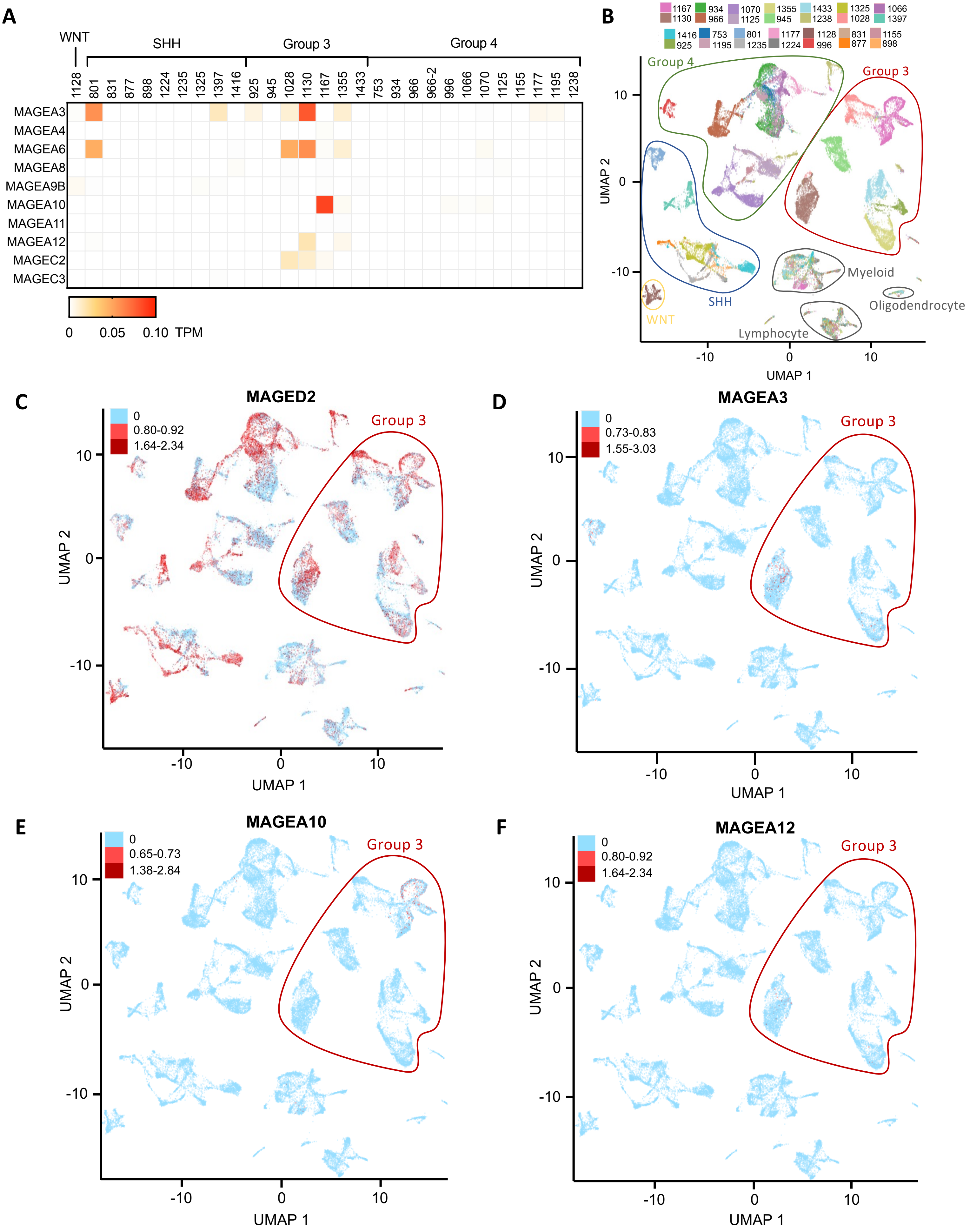


**Supp. Figure 2**

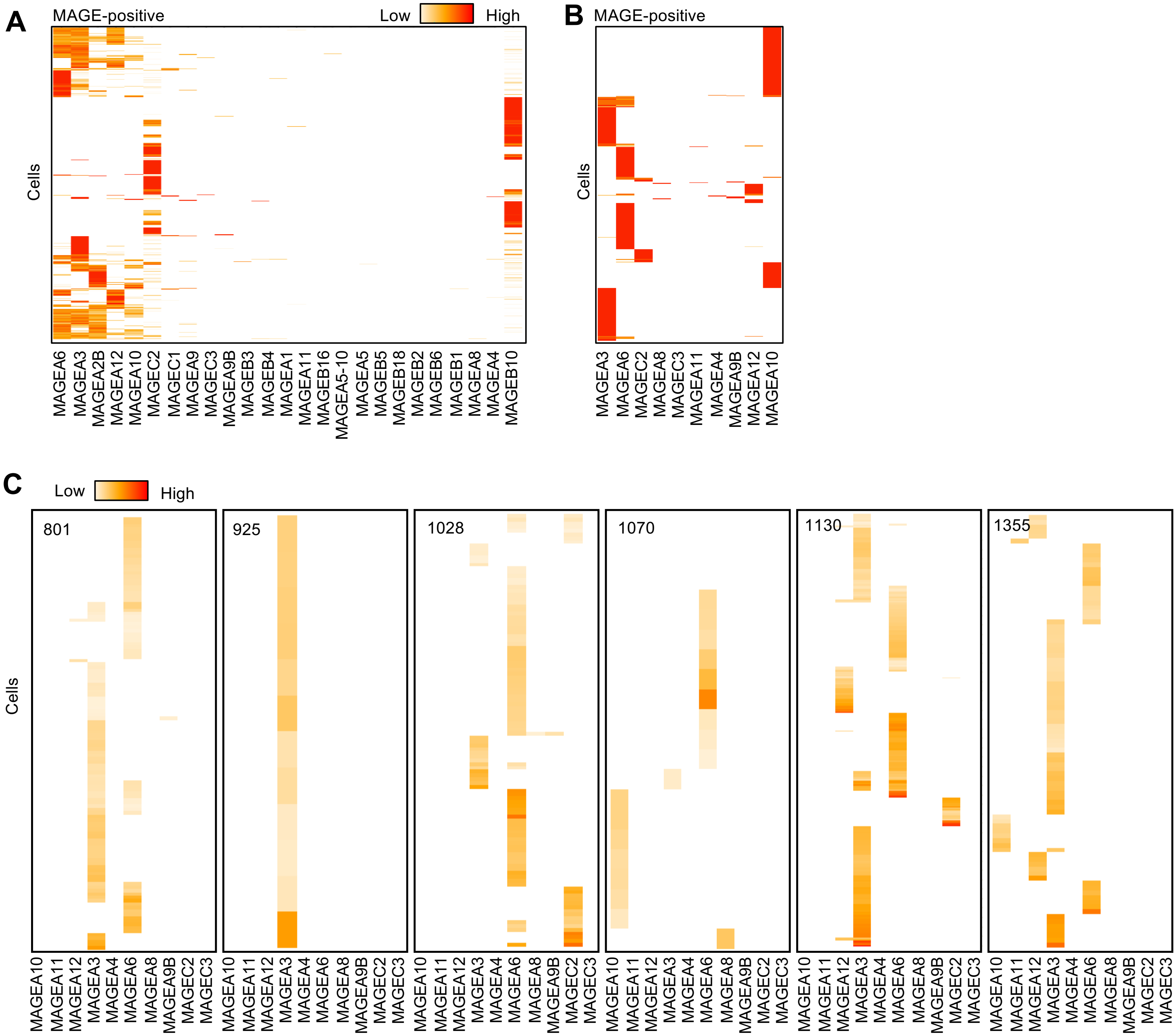


**Supp. Figure 3**

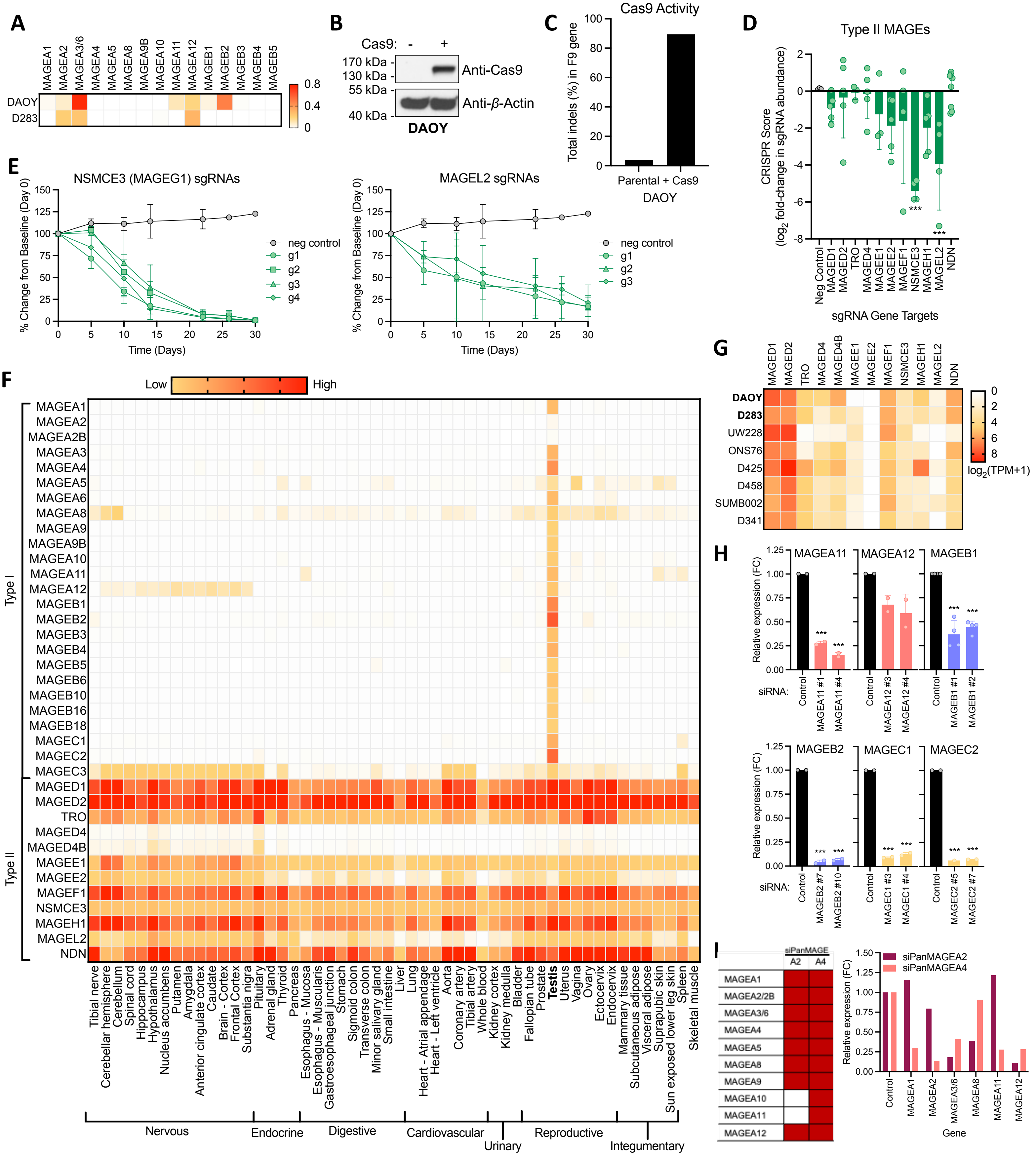


**Supp. Figure 4**

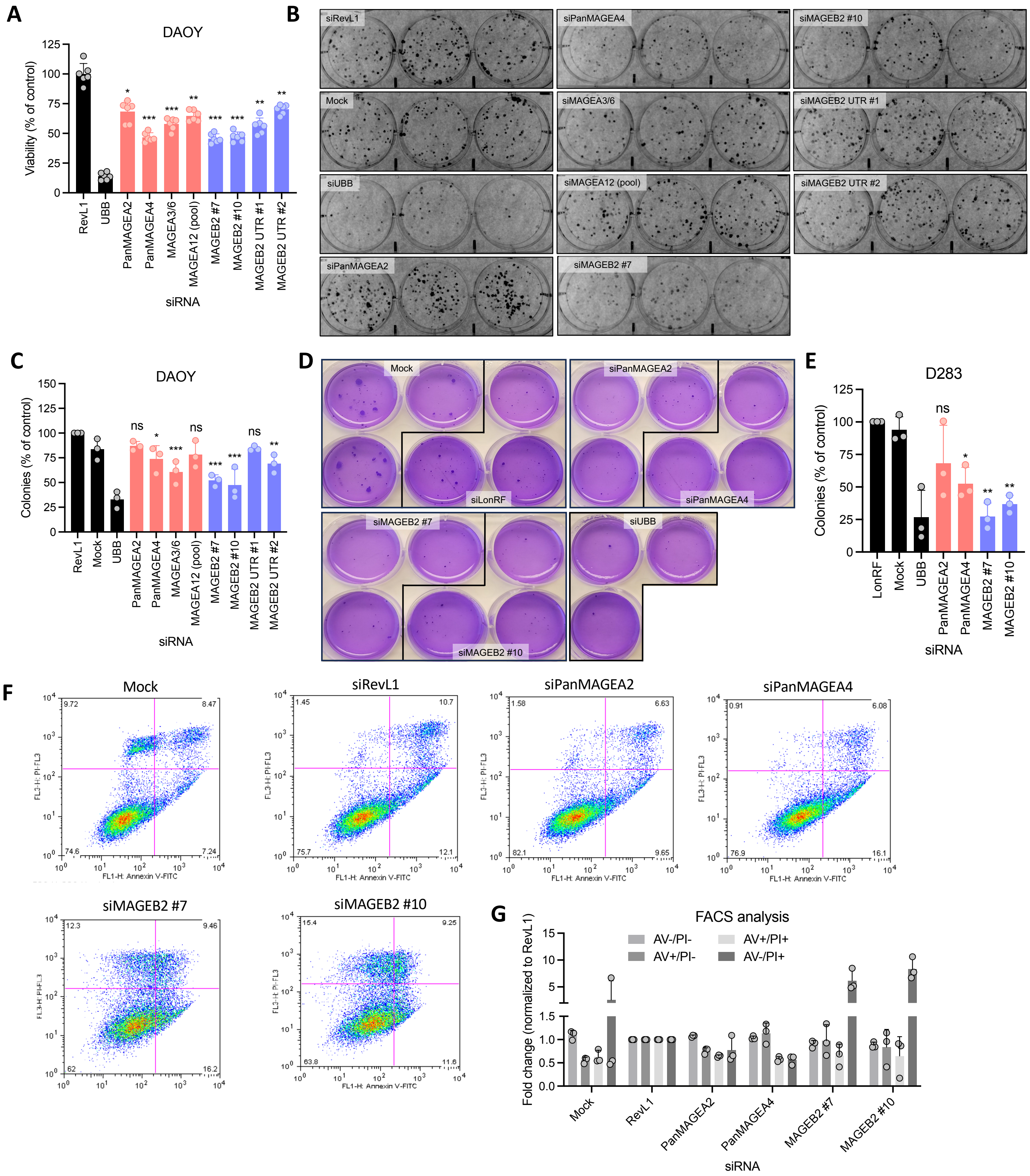


**Supp. Figure 5**

Table S4. Percentage of group 3-MB cells expressing each type I MAGE

| Type I MAGEs | MUV11 | SJ17 | SJ917 | SJ617 | MUV29 | BCH1205 | MUV34 | BCH825 |
|---|---|---|---|---|---|---|---|---|
| MAGEA1 | 3.0 | 0.0 | 0.0 | 1.8 | 0.3 | 0.6 | 0.0 | 0.0 |
| MAGEA2 | 20.9 | 1.8 | 0.0 | 27.6 | 2.9 | 36.5 | 3.4 | 1.0 |
| MAGEA2B | 32.5 | 5.1 | 0.0 | 37.8 | 8.0 | 58.9 | 8.2 | 2.4 |
| MAGEA3 | 57.5 | 1.8 | 0.3 | 64.6 | 12.7 | 79.8 | 10.9 | 0.3 |
| MAGEA4 | 0.2 | 0.0 | 0.3 | 0.9 | 0.6 | 0.0 | 0.0 | 0.3 |
| MAGEA5 | 0.2 | 0.0 | 0.0 | 0.6 | 0.0 | 0.6 | 0.0 | 0.3 |
| MAGEA6 | 51.3 | 1.8 | 0.0 | 58.9 | 8.9 | 67.2 | 7.1 | 0.7 |
| MAGEA8 | 0.2 | 0.0 | 0.3 | 0.0 | 0.3 | 0.3 | 0.0 | 0.0 |
| MAGEA9 | 0.9 | 0.3 | 0.0 | 0.0 | 1.0 | 0.6 | 0.7 | 0.0 |
| MAGEA9B | 0.2 | 0.0 | 0.0 | 0.0 | 0.0 | 0.3 | 0.0 | 0.3 |
| MAGEA10 | 23.9 | 23.1 | 26.4 | 38.7 | 32.8 | 41.7 | 19.7 | 48.6 |
| MAGEA11 | 0.2 | 0.0 | 0.0 | 0.0 | 0.0 | 0.9 | 0.0 | 0.0 |
| MAGEA12 | 42.9 | 0.0 | 0.0 | 38.4 | 4.8 | 22.4 | 0.0 | 1.0 |
| MAGEB1 | 0.0 | 0.0 | 0.0 | 0.0 | 0.0 | 0.3 | 0.0 | 0.0 |
| MAGEB2 | 0.0 | 0.0 | 0.0 | 0.0 | 0.0 | 0.0 | 0.0 | 0.0 |
| MAGEB3 | 0.2 | 0.0 | 0.3 | 0.0 | 0.3 | 0.9 | 0.0 | 0.0 |
| MAGEB4 | 0.5 | 0.0 | 0.0 | 0.0 | 0.3 | 0.9 | 0.0 | 0.0 |
| MAGEB5 | 0.0 | 0.0 | 0.0 | 0.0 | 0.0 | 0.0 | 0.0 | 0.0 |
| MAGEB6 | 0.2 | 0.0 | 0.0 | 0.3 | 0.0 | 0.0 | 0.0 | 0.0 |
| MAGEB10 | 20.6 | 30.5 | 32.4 | 38.4 | 31.5 | 46.0 | 23.1 | 60.9 |
| MAGEB16 | 0.5 | 0.0 | 0.0 | 0.0 | 0.0 | 0.6 | 0.0 | 0.3 |
| MAGEB18 | 0.0 | 0.0 | 0.0 | 0.0 | 0.0 | 0.0 | 0.0 | 0.0 |
| MAGEC1 | 0.0 | 0.0 | 0.0 | 3.6 | 1.9 | 10.4 | 0.0 | 0.3 |
| MAGEC2 | 0.0 | 0.0 | 0.0 | 0.0 | 3.2 | 7.1 | 0.7 | 0.7 |
| MAGEC3 | 0.2 | 0.0 | 1.2 | 0.0 | 0.3 | 0.3 | 0.0 | 0.7 |

**Table S5. Percentage of group 3-MB cells expressing # of type I MAGEs**

| # of MAGEs | MUV11 | SJ17 | SJ917 | SJ617 | MUV29 | BCH1205 | MUV34 | BCH825 |
|---|---|---|---|---|---|---|---|---|
| 0 | 12.3 | 49.7 | 51.6 | 3.9 | 39.5 | 3.1 | 54.1 | 27.6 |
| 1 | 18.6 | 38.6 | 35.7 | 12.6 | 32.8 | 6.1 | 26.9 | 45.2 |
| 2 | 20.6 | 9.9 | 12.3 | 23.7 | 16.2 | 13.8 | 13.9 | 26.2 |
| 3 | 18.1 | 1.2 | 0.3 | 21.3 | 4.5 | 20.9 | 2.7 | 1.0 |
| 4 | 14.9 | 0.6 | 0.0 | 18.3 | 4.5 | 23.9 | 1.7 | 0.0 |
| 5 | 11.8 | 0.0 | 0.0 | 12.3 | 2.2 | 15.6 | 0.7 | 0.0 |
| > 5 | 3.7 | 0.0 | 0.0 | 7.8 | 0.3 | 16.6 | 0.0 | 0.0 |

**Supp. Table 5**

| Sample | Molecular Subtype (by our qPCR data) | Supporting Data Confirm? | St Jude, MYC/MYCN amp, or IHC | Gender | Age at Dx | Clinical Risk Stratification at Dx | Intracranial spread/CSF+/Drop_ mets/Mets? | Is this tissue from a met? | Survival from Dx | Histologic Subtype | Histology Features and Comments |
|---|---|---|---|---|---|---|---|---|---|---|---|
| MT_2.(1+2) | G3/4 | | | M | 3 y | High Risk | Y | N | 18 days | Large Cell Variant | Large Cell Variant |
| MT_4.1 | SHH | | | M | 6 y | Avg | N | N | 4 y | Classic | Classic medulloblastoma. Although there are some features resembling a nodular / desmoplastic medulloblastoma |
| MT_5.(1+2) | SHH | | | M | 1 y | Infant (<3) | N | N | + | Classic | Suggests desmoplastic medulloblastoma; however, the nodularity in this case is not especially well developed |
| MT_7.(1+2) | SHH | | | F | 9 m | Infant (<3) | N | N | + | Classic | Classic |
| MT_13.(1+2) | G3/4 | | | M | 6 y | High Risk | Y | N | + | Classic | Although foci of pleomorphic morphology and larger cells are noted, this is not present ito a sufficient degree to warrant the diagnosis of anaplastic medulloblastoma |
| MT_14.(1+2) | G3/4 | | | M | 6 y | High Risk | Y | N | + | Classic | Classic |
| MT_15.(1+2) | G3/4 | | | F | 7 y | High Risk | Y | N | + | Anaplastic variant | Anaplastic variant |
| MT_16.(1+2) | G3/4 | | | F | 2 y | Infant (<3) | N | N | + | Extensive nodularity | Extensive nodularity |
| MT_17.1 | SHH | | | F | 3 y | Avg | N | N | + | extensive nodularity | Desmoplastic with extensive nodularity |
| MT_18.1 | SHH | | | M | 15 y | Avg | N | N | Unknown | nodularity | Desmoplastic with focal nodularity |
| MT_19.1 | SHH | | | M | 17 y | Avg | N | N | + | Classic | Classic |
| MT_21.4 | SHH | | | M | 14 y | High Risk | Y | N | 4 y | Classic | Classic |
| MT_22.1 | G3/4 | | | M | 11 y | High Risk | Y | N | + | Classic | Classic with Features of anaplasia |
| MT_23.(1+2) | --- | | | F | 3 y | High Risk | Y | N | 10 mo | Classic | Classic |
| MT_24.1 | G3/4 | | | F | 10 y | High Risk | Unknown | N | + | Classic | Classic with features of possible focal nodular phenotype, but no desmoplasia |
| MT_25.1 | G3/4 | | | M | 4 y | High Risk | Y | N | + | melanotic differentiation | Medulloblastoma with melanotic differentiation |
| MT_26.1 | SHH | | | M | 13 y | High Risk | Y | N | + | Desmoplastic/nodular type | Desmoplastic/nodular type |
| MT_27.1 | G3/4 | | | M | 8 y | High Risk | Y | N | + | Classic | Classic, with focal features of anaplasia |
| MT_29.1 | SHH | | St. Jude - unable to classify by IHC | M | 23 week | Infant (<3) | N | N | + | Desmoplastic/nodular type | Desmoplastic/nodular type |
| MT_30.(1+2) | G3/4 | | | M | 8 y | High Risk | Y | N | 3.8 y | Classic | Classic |
| MT_31.2 | G3/4 | ✶ | St. Jude - MycN amp, G3/G4 | M | 15 y | Avg | N | N | 1.3 y | Desmoplastic/nodular type | Desmoplastic/nodular type |
| MT_32.2 | G3/4 | | | M | 6 y | Avg | N | N | + | Classic | Classic |
| MT_33.1 | G3/4 | | | M | 3 y | Avg | N | N | + | Classic | Classic |
| MT_34.1 | WNT | ✶ | CMC IHC myc+ | F | 5 y | Avg | N | N | Unknown | Large Cell Variant | Large Cell Variant |
| MT_35.1 | G3/4 | | | M | 6 y | Avg | N | N | + | moderate anaplasia | Medulloblastoma with moderate anaplasia |
| MT_36.1 | G3/4 | | | F | 5 y | Avg | N | N | + | Classic | Classic |
| MT_37.2 | G3/4 | ✶ | CMC IHC myc neg | F | 4 y | Avg | N | N | + | with focal desmoplasia | Large cell/Anaplastic type with focal desmoplasia |
| MT_38.1 | G3/4 | | | F | 4 y | High Risk | Y | N | + | Anaplastic variant | Anaplastic variant |
| MT_39.1 | SHH | ✶ | St Jude - SHH (YAP and GAB Positive, myc and mycN amp) | M | 14 mo | Infant (<3) | Y | N | 1 y | Classic | Medulloblastoma with anaplastic and nodular/desmoplastic features |
| MT_40.1 | SHH | ✶ | myc and mycN amp) | F | 17 mo | Infant (<3) | Unknown | N | 1.2 y | Large cell/Anaplastic type | Large cell/Anaplastic type |
| MT_41.1 | G3/4 | | | M | 11 y | | N | N | 1 y | Large cell/Anaplastic | Large cell/Anaplastic |
| MT_42.1 | SHH | ✶ | CMC IHC myc neg | F | 2 y | Infant (<3) | Y | N | 11 mo | Desmoplastic/nodular type | Desmoplastic/nodular type |
| MT_43.2 | G3/4 | ✶ | St Jude - G3/G4, no myc/mycN amp | M | 11 mo | Infant (<3) | N | N | + | Classic | Classic |
| MT_44.1 | G3/4 | ✶ | and GAB negative | M | 4 y | High Risk | Y | N | + | Classic | Classic |

+ = Patient alive at time of recruitment

Table S2 - MAGE_gRNA_library

| sgRNA | sequence | target |
|---|---|---|
| **SNP116.mMAGEA4.g30** | TTTGGCTCCGAGCACTGCCT | mMAGEA4 |
| **SNP116.mMAGEA4.g31** | GGCTCCGAGCACTGCCTTGG | mMAGEA4 |
| **SNP116.mMAGEA4.g35** | CTGCAGTAATGGGTGTTATC | mMAGEA4 |
| **SNP116.mMAGEA4.g4** | AGGCAGTGCTCGGAGCCAAA | mMAGEA4 |
| **SNP117.mMAGEA5.g20** | CTCCTCCTCGGATGTCTCCA | mMAGEA5 |
| **SNP117.mMAGEA5.g21** | CGGATGTCTCCATGGTCTCC | mMAGEA5 |
| **SNP117.mMAGEA5.g22** | GGTCATTGTCTAATTCCTGT | mMAGEA5 |
| **SNP119.mMAGEA7PS.g12** | ACCATAGACCTTATTTGAGG | mMAGEA7PS |
| **SNP138.mMAGEB16.g18** | ACATGGCTCCTCAACATCGA | mMAGEB16 |
| **SNP138.mMAGEB16.g9** | CCAAGTCTCCAAGGATGTGG | mMAGEB16 |
| **SNP140.mGM5072.g34** | GGCTGAGCACCCTGGAGTTT | mGM5072 |
| **SNP143.m4930550L24RIK.g18** | GCTTGATGCAGACGCTTGGC | m4930550L24RIK |
| **SNP145.mMAGED2.g19** | TTGAGACCTCTGTCGCCTTT | mMAGED2 |
| **SNP81.hMAGEA8.g19** | CAGAATCAGTCACCTCCTCA | hMAGEA8 |
| **SNP87.hMAGEB1.g29** | GTGAGCAACCTTGAGACCCT | hMAGEB1 |
| **SNP88.hMAGEB2.g1** | TGCCCGTGAGAAACGCCGCA | hMAGEB2 |
| **SNP88.hMAGEB2.g27** | AGCTTACTCTTCTGACCACG | hMAGEB2 |
| **SNP96.hMAGEB18.g13** | GATTCTCACAACGAGCCTGG | hMAGEB18 |
| **SNP96.hMAGEB18.g4** | CGTTGTGAGAATCAGGATCT | hMAGEB18 |
| **SNP97.hMAGEC1.g15** | AGACTCGGCATCCCAGCAGT | hMAGEC1 |
| **SNP98.hMAGEC2.g11** | TTCCTCCTCTTCCTCATCTG | hMAGEC2 |
| **SNP98.hMAGEC2.g5** | TCCCACAGATGAGGAAGAGG | hMAGEC2 |
| **SM76_negative_eGFP.mneg.g5** | AGCACTGCACGCCGTAGGTC | mneg |
| **SM76_negative_eGFP.mneg.g6** | GCACTGCACGCCGTAGGTCA | mneg |
| **SM76_negative_human.hneg.g22** | TAAGAGGGCGAAGGAGTCAT | hneg |
| **SM76_negative_human.hneg.g27** | ATGGAAGGCTCGCCTATCCT | hneg |
| **SM76_negative_human.hneg.g24** | TAGGCGAGCCTTCCAAGGAT | hneg |
| **SM76_negative_human.hneg.g25** | AAGGATAGGCGAGCCTTCCA | hneg |
| **SM76_negative_human.hneg.g15** | GCACAGGGGCTTGTCAATCG | hneg |

| | | |
|---|---|---|
| **SM76_negative_human.hneg.g34** | CTGCCTATGGCCTAACTCCA | hneg |
| **SM76_negative_human.hneg.g9** | GCTTGTGGATGTTGCGGAAG | hneg |
| **SM76_negative_mCherry.mneg.g19** | AGTAGTCGGGGATGTCGGCG | mneg |
| **SM76_negative_mCherry.mneg.g17** | CAAGTAGTCGGGGATGTCGG | mneg |
| **SM76_negative_mouse.mneg.g15** | GCACAGGGGCTTGTCAATCG | mneg |
| **SNP100.hMAGED1.g15** | TCTGGCAGACGCCATTGGCT | hMAGED1 |
| **SNP100.hMAGED1.g2** | ATGTTGAAGAGAACAGCAGT | hMAGED1 |
| **SNP100.hMAGED1.g26** | CATTGGGGGGTGCGCCAGGT | hMAGED1 |
| **SNP100.hMAGED1.g29** | TTCTGAGTGGTCACTGGAAC | hMAGED1 |
| **SNP100.hMAGED1.g7** | GGATCAGAGGCGGGCCCCAC | hMAGED1 |
| **SNP101.hMAGED2.g1** | GTTGACAGTGACCCAGAATG | hMAGED2 |
| **SNP101.hMAGED2.g11** | TTTGAGGCCTTCGGTGTCTC | hMAGED2 |
| **SNP101.hMAGED2.g12** | TTGAGGCCTTCGGTGTCTCT | hMAGED2 |
| **SNP101.hMAGED2.g2** | GACAGTGACCCAGAATGTGG | hMAGED2 |
| **SNP101.hMAGED2.g6** | GAAGGCCTCAAAGGCACTGG | hMAGED2 |
| **SNP102.hTRO.g1** | GACCAAGGACACACCCAAGC | hTRO |
| **SNP102.hTRO.g10** | ACCCAGCTTGGGTGTGTCCT | hTRO |
| **SNP102.hTRO.g4** | CTGAGTGTCATTTTTATGAA | hTRO |
| **SNP102.hTRO.g5** | CATTTTTATGAATGGCAACA | hTRO |
| **SNP102.hTRO.g7** | TGACACTCAGAATCACCATG | hTRO |
| **SNP103.hMAGED4.g12** | TATGAAGAATTCGGTGCGTT | hMAGED4 |
| **SNP103.hMAGED4.g17** | CTCAGAGCCTTCGGAAGCTT | hMAGED4 |
| **SNP103.hMAGED4.g19** | TCCAGGCCTTCGCATCTTAT | hMAGED4 |
| **SNP103.hMAGED4.g6** | ATGAGGGTAGCGACGAAGTC | hMAGED4 |
| **SNP103.hMAGED4.g7** | TGAGGGTAGCGACGAAGTCG | hMAGED4 |
| **SNP105.hMAGEE1.g1** | CCGCCGCCGCCGCGTTGCAA | hMAGEE1 |
| **SNP105.hMAGEE1.g35** | GCGCAGTAGCCTTTGCAACG | hMAGEE1 |
| **SNP105.hMAGEE1.g36** | CAGTAGCCTTTGCAACGCGG | hMAGEE1 |
| **SNP105.hMAGEE1.g37** | TAGCCTTTGCAACGCGGCGG | hMAGEE1 |
| **SNP106.hMAGEE2.g16** | GGCATCAACGACTAGCATGG | hMAGEE2 |
| **SNP106.hMAGEE2.g23** | CAGTGATCTCTGCGCTACAG | hMAGEE2 |
| **SNP106.hMAGEE2.g24** | CGCTACAGTGGCGTGCATTC | hMAGEE2 |

| | | |
|---|---|---|
| **SNP106.hMAGEE2.g3** | GCAGATTACGGCGACGGCAG | hMAGEE2 |
| **SNP106.hMAGEE2.g4** | ATACAAGCTACTAACGCCTC | hMAGEE2 |
| **SNP107.hMAGEF1.g14** | GAGAAGGATGGCGGCCATGA | hMAGEF1 |
| **SNP107.hMAGEF1.g44** | CCTGCGAGGCGGTCGGGGCC | hMAGEF1 |
| **SNP107.hMAGEF1.g52** | ACCGGGAGCCCCCTGCTCTC | hMAGEF1 |
| **SNP107.hMAGEF1.g6** | CAGGGGGCTCCCGGTCCCGC | hMAGEF1 |
| **SNP107.hMAGEF1.g8** | CTCCCGGTCCCGCAGGCCGA | hMAGEF1 |
| **SNP108.hNSMCE3.g1** | TGTTGCAAAAACCGAGGAAC | hNSMCE3 |
| **SNP108.hNSMCE3.g15** | CGGAAACCCCGGGGCTTCGC | hNSMCE3 |
| **SNP108.hNSMCE3.g2** | GTTGCAAAAACCGAGGAACC | hNSMCE3 |
| **SNP108.hNSMCE3.g3** | TTGCAAAAACCGAGGAACCG | hNSMCE3 |
| **SNP109.hMAGEH1.g2** | CCGCAACAATCGCAAAATCC | hMAGEH1 |
| **SNP109.hMAGEH1.g22** | CTTCTGCGGCTCTCGCATTA | hMAGEH1 |
| **SNP109.hMAGEH1.g23** | CTGCGGCTCTCGCATTACGG | hMAGEH1 |
| **SNP109.hMAGEH1.g24** | CGCCGACTCTTTCGTCCCCG | hMAGEH1 |
| **SNP109.hMAGEH1.g4** | AGAGGCCTCCGAGACCCCTA | hMAGEH1 |
| **SNP110.hMAGEL2.g1** | GTCGCAGCTAAGTAAGAATC | hMAGEL2 |
| **SNP110.hMAGEL2.g2** | TCGCAGCTAAGTAAGAATCT | hMAGEL2 |
| **SNP110.hMAGEL2.g3** | TCTGGGTGACTCGAGTCCTC | hMAGEL2 |
| **SNP110.hMAGEL2.g37** | GCTATAGACAGGCGGCTTCG | hMAGEL2 |
| **SNP110.hMAGEL2.g5** | TGACTCGAGTCCTCCGGCGG | hMAGEL2 |
| **SNP111.hNDN.g10** | CAGCCCTGGGGTTTCGGAGG | hNDN |
| **SNP111.hNDN.g13** | GAGCCCTCCTCTAGGCCCGA | hNDN |
| **SNP111.hNDN.g18** | GGAGCGGCCGTCGGGCCTAG | hNDN |
| **SNP111.hNDN.g19** | GCGGCCGTCGGGCCTAGAGG | hNDN |
| **SNP111.hNDN.g27** | GGAACCCCCTCCGAAACCCC | hNDN |
| **SNP113.mMAGEA1.g26** | AAGTATTGGTAGAGAGTATG | mMAGEA1 |
| **SNP113.mMAGEA1.g27** | CTACCCTCTGATCTTTAGTG | mMAGEA1 |
| **SNP113.mMAGEA1.g35** | GCCCTGGGGATCACCTATGA | mMAGEA1 |
| **SNP113.mMAGEA1.g49** | GAGGCCTCACTAAAGATCAG | mMAGEA1 |
| **SNP113.mMAGEA1.g5** | GGAAACCATGGAGACGTCAG | mMAGEA1 |
| **SNP113.mMAGEA1.g50** | AGGCCTCACTAAAGATCAGA | mMAGEA1 |

| | | |
|---|---|---|
| **SNP113.mMAGEA1.g56** | TTGATGGAAGCTTCCTCAGA | mMAGEA1 |
| **SNP113.mMAGEA1.g6** | AACCATGGAGACGTCAGAGG | mMAGEA1 |
| **SNP113.mMAGEA1.g70** | TGAGGACTCTGGGGAGGACT | mMAGEA1 |
| **SNP113.mMAGEA1.g74** | CTCCTCCTCTGACGTCTCCA | mMAGEA1 |
| **SNP113.mMAGEA1.g75** | CTGACGTCTCCATGGTTTCC | mMAGEA1 |
| **SNP113.mMAGEA1.g83** | GCTGCAGTATTGGGTGTTAC | mMAGEA1 |
| **SNP114.mMAGEA2.g1** | CAGCCTCCAAGAGAGTGCTC | mMAGEA2 |
| **SNP114.mMAGEA2.g18** | ATGGAGGCCAGTGCCACACA | mMAGEA2 |
| **SNP114.mMAGEA2.g20** | CAGGGAGAGTAGGCTCTCTG | mMAGEA2 |
| **SNP114.mMAGEA2.g22** | GTAGGCTCTCTGAGGACTCT | mMAGEA2 |
| **SNP114.mMAGEA2.g24** | GCTCTCTGAGGACTCTGGGG | mMAGEA2 |
| **SNP114.mMAGEA2.g35** | TCTCTTGGAGGCTGCAGTAT | mMAGEA2 |
| **SNP114.mMAGEA2.g36** | CTCTTGGAGGCTGCAGTATT | mMAGEA2 |
| **SNP114.mMAGEA2.g37** | GGCTGCAGTATTGGGTGTTA | mMAGEA2 |
| **SNP114.mMAGEA2.g38** | GCTGCAGTATTGGGTGTTAT | mMAGEA2 |
| **SNP115.mMAGEA3.g1** | CAACCTCGAAGAGAGTGCTC | mMAGEA3 |
| **SNP115.mMAGEA3.g11** | TAGGCTCTCTGAGGACTCTG | mMAGEA3 |
| **SNP115.mMAGEA3.g16** | ATACACTTTATTTGAGGTGG | mMAGEA3 |
| **SNP115.mMAGEA3.g2** | AGAGAGTGCTCAGGCCCAAC | mMAGEA3 |
| **SNP115.mMAGEA3.g21** | GGGCCTGAGCACTCTCTTCG | mMAGEA3 |
| **SNP115.mMAGEA3.g22** | TCTCTTCGAGGTTGCAGTAT | mMAGEA3 |
| **SNP115.mMAGEA3.g23** | CTCTTCGAGGTTGCAGTATT | mMAGEA3 |
| **SNP115.mMAGEA3.g25** | GTTGCAGTATTGGGTGTTAT | mMAGEA3 |
| **SNP115.mMAGEA3.g8** | ACCACCTCAAATAAAGTGTA | mMAGEA3 |
| **SNP115.mMAGEA3.g9** | AGTAGGCTCTCTGAGGACTC | mMAGEA3 |
| **SNP116.mMAGEA4.g14** | GAACAGAGGCTAGTATCACA | mMAGEA4 |
| **SNP116.mMAGEA4.g23** | GCTCTTTGAGGACTCTGGGG | mMAGEA4 |
| **SNP117.mMAGEA5.g23** | GTCATTGTCTAATTCCTGTT | mMAGEA5 |
| **SNP117.mMAGEA5.g25** | GGGCCTGAGCACTCTCTTGG | mMAGEA5 |
| **SNP117.mMAGEA5.g3** | ACAGGAATTAGACAATGACC | mMAGEA5 |
| **SNP117.mMAGEA5.g4** | AGACAATGACCAGGAGACCA | mMAGEA5 |
| **SNP117.mMAGEA5.g5** | GGAGACCATGGAGACATCCG | mMAGEA5 |

| | | |
|---|---|---|
| **SNP117.mMAGEA5.g6** | GACCATGGAGACATCCGAGG | mMAGEA5 |
| **SNP117.mMAGEA5.g7** | CATGGAGACATCCGAGGAGG | mMAGEA5 |
| **SNP118.mMAGEA6.g16** | ACTGCCATACACTTTATTTG | mMAGEA6 |
| **SNP118.mMAGEA6.g17** | GCCATACACTTTATTTGAGG | mMAGEA6 |
| **SNP118.mMAGEA6.g2** | AGAGAGTGCTCAGGCTCAAC | mMAGEA6 |
| **SNP118.mMAGEA6.g21** | GAGCCTGAGCACTCTCTTCG | mMAGEA6 |
| **SNP118.mMAGEA6.g22** | TCTCTTCGAGGTTGCAGTAT | mMAGEA6 |
| **SNP118.mMAGEA6.g23** | CTCTTCGAGGTTGCAGTATT | mMAGEA6 |
| **SNP118.mMAGEA6.g24** | GGTTGCAGTATTGGGTGTTA | mMAGEA6 |
| **SNP118.mMAGEA6.g3** | ACAGGAATCAGACAATGACC | mMAGEA6 |
| **SNP118.mMAGEA6.g9** | GAGAGCCTACTCTCCCTGTG | mMAGEA6 |
| **SNP119.mMAGEA7PS.g1** | AGAGAGTGCTCAGTCCCAAC | mMAGEA7PS |
| **SNP119.mMAGEA7PS.g10** | TGAGGACTCTGGGGAGGATT | mMAGEA7PS |
| **SNP119.mMAGEA7PS.g11** | ACTACCATAGACCTTATTTG | mMAGEA7PS |
| **SNP119.mMAGEA7PS.g15** | CTGATGTGTCCATGGTAGCC | mMAGEA7PS |
| **SNP119.mMAGEA7PS.g18** | GTTGGGACTGAGCACTCTCT | mMAGEA7PS |
| **SNP119.mMAGEA7PS.g19** | GGGACTGAGCACTCTCTTGG | mMAGEA7PS |
| **SNP119.mMAGEA7PS.g3** | AGATAATGACCAGGCTACCA | mMAGEA7PS |
| **SNP119.mMAGEA7PS.g7** | AGATACCACCACCTCAAATA | mMAGEA7PS |
| **SNP119.mMAGEA7PS.g9** | TCAAATAAGGTCTATGGTAG | mMAGEA7PS |
| **SNP120.mMAGEA8.g1** | CAACCTCCAAGAGAGTGCTC | mMAGEA8 |
| **SNP120.mMAGEA8.g15** | CTCCTCCTGTGATGTCACCA | mMAGEA8 |
| **SNP120.mMAGEA8.g16** | GTGATGTCACCATGGTCTCC | mMAGEA8 |
| **SNP120.mMAGEA8.g19** | GTTGGGCCTGAGCACTCTCT | mMAGEA8 |
| **SNP120.mMAGEA8.g21** | TCTCTTGGAGGTTGCAGTAT | mMAGEA8 |
| **SNP120.mMAGEA8.g5** | GGAGACCATGGTGACATCAC | mMAGEA8 |
| **SNP120.mMAGEA8.g6** | GACCATGGTGACATCACAGG | mMAGEA8 |
| **SNP120.mMAGEA8.g7** | CATGGTGACATCACAGGAGG | mMAGEA8 |
| **SNP121.mMAGEA9PS.g1** | TCCAATATCCCCAACTGATG | mMAGEA9PS |
| **SNP121.mMAGEA9PS.g18** | TGACAGAGGCCTCATCAGTT | mMAGEA9PS |
| **SNP121.mMAGEA9PS.g22** | TGGAATGTTGGCTATAACAA | mMAGEA9PS |
| **SNP121.mMAGEA9PS.g23** | GGAATGTTGGCTATAACAAT | mMAGEA9PS |

| | | |
|---|---|---|
| **SNP121.mMAGEA9PS.g24** | ATGTTGGCTATAACAATGGG | mMAGEA9PS |
| **SNP122.mMAGEA10.g1** | TGCCTAGACCCCGGAAGCGT | mMAGEA10 |
| **SNP122.mMAGEA10.g23** | CCATACATCTCCTACGCTTC | mMAGEA10 |
| **SNP122.mMAGEA10.g24** | CATACATCTCCTACGCTTCC | mMAGEA10 |
| **SNP122.mMAGEA10.g25** | ATACATCTCCTACGCTTCCG | mMAGEA10 |
| **SNP122.mMAGEA10.g26** | CTCCTACGCTTCCGGGGTCT | mMAGEA10 |
| **SNP123.m1700080O16RIK.g1** | TGGATCCTCCCGACAACAGC | m1700080O16RIK |
| **SNP123.m1700080O16RIK.g2** | CAACAGCCGGAATGTCAACC | m1700080O16RIK |
| **SNP123.m1700080O16RIK.g30** | TTTCAGAATGAGTGCTGTCC | m1700080O16RIK |
| **SNP123.m1700080O16RIK.g33** | TTGACATTCCGGCTGTTGTC | m1700080O16RIK |
| **SNP123.m1700080O16RIK.g34** | ACATTCCGGCTGTTGTCGGG | m1700080O16RIK |
| **SNP124.m3830417A13RIK.g18** | GCCCCAGGGGATGTGCCTGC | m3830417A13RIK |
| **SNP124.m3830417A13RIK.g25** | ACTGCCATGGGTTCTATTTC | m3830417A13RIK |
| **SNP124.m3830417A13RIK.g52** | TAGGGTCCTGGGTACTCCCG | m3830417A13RIK |
| **SNP124.m3830417A13RIK.g58** | TGTCCCATAGGTTGAAGTTC | m3830417A13RIK |
| **SNP124.m3830417A13RIK.g8** | CAAACTGAAGAAAGGGGCCC | m3830417A13RIK |
| **SNP125.m4933402E13RIK.g14** | AGAGGAAGATAACGTCGAGA | m4933402E13RIK |
| **SNP125.m4933402E13RIK.g2** | CCACGCCTCAGTCTGGAAGA | m4933402E13RIK |
| **SNP125.m4933402E13RIK.g21** | TCATCGTCATCAGTTTCCGT | m4933402E13RIK |
| **SNP125.m4933402E13RIK.g25** | CTTCTTCCAGACTGAGGCGT | m4933402E13RIK |
| **SNP125.m4933402E13RIK.g3** | TGTCAACTTCCAGAATCCAA | m4933402E13RIK |
| **SNP126.mMAGEB1.g14** | TAGGCATGTTGACTGTGCTC | mMAGEB1 |
| **SNP126.mMAGEB1.g2** | CCAGGTCTCCATTAAGTCCA | mMAGEB1 |
| **SNP126.mMAGEB1.g20** | GTAAGTACCTCTGTACTACC | mMAGEB1 |
| **SNP126.mMAGEB1.g22** | AGAGAATACCTTGGACTTAA | mMAGEB1 |
| **SNP126.mMAGEB1.g3** | AGTCCAAGGTATTCTCTACC | mMAGEB1 |
| **SNP126.mMAGEB1.g4** | TTCTCTACCTGGTAGTACAG | mMAGEB1 |
| **SNP126.mMAGEB1.g6** | TGAGCACAGTCAACATGCCT | mMAGEB1 |
| **SNP126.mMAGEB1.g7** | GAGCACAGTCAACATGCCTA | mMAGEB1 |
| **SNP126.mMAGEB1.g8** | AGCACAGTCAACATGCCTAG | mMAGEB1 |
| **SNP127.mMAGEB2.g1** | CAAAACGACAGCAGTCACGC | mMAGEB2 |
| **SNP127.mMAGEB2.g15** | CCCTGAGGAGCAGAACCACC | mMAGEB2 |

| | | |
|---|---|---|
| **SNP127.mMAGEB2.g2** | AAAACGACAGCAGTCACGCA | mMAGEB2 |
| **SNP127.mMAGEB2.g24** | TGACTGCTGTCGTTTTGCAC | mMAGEB2 |
| **SNP127.mMAGEB2.g26** | CTGTCGTTTTGCACGGGAGC | mMAGEB2 |
| **SNP127.mMAGEB2.g8** | CTCCTGTTGACCAGAGTGCT | mMAGEB2 |
| **SNP128.mMAGEB3.g1** | CTAAACGACAGCAGTCACGC | mMAGEB3 |
| **SNP128.mMAGEB3.g11** | GTGACTGCTGTCGTTTAGCA | mMAGEB3 |
| **SNP128.mMAGEB3.g12** | TGACTGCTGTCGTTTAGCAC | mMAGEB3 |
| **SNP128.mMAGEB3.g2** | TAAACGACAGCAGTCACGCA | mMAGEB3 |
| **SNP128.mMAGEB3.g7** | TCTCCTGTTGACCTGAGTGC | mMAGEB3 |
| **SNP129.mGM44.g14** | GAGCATAATTTCTGGCACGG | mGM44 |
| **SNP129.mGM44.g15** | GGCACGGTGGCGTCTCTCCC | mGM44 |
| **SNP129.mGM44.g2** | AAAGAGTAAGCTAAATGCCA | mGM44 |
| **SNP129.mGM44.g3** | CCGTGCCAGAAATTATGCTC | mGM44 |
| **SNP130.mGM16441.g12** | GGTCCCCATAGCAGGGGCAA | mGM16441 |
| **SNP130.mGM16441.g15** | GAGACCGTTGCAGCTACCTC | mGM16441 |
| **SNP130.mGM16441.g16** | AGACCGTTGCAGCTACCTCA | mGM16441 |
| **SNP130.mGM16441.g8** | GGTACACAGGTCCCCATAGC | mGM16441 |
| **SNP130.mGM16441.g9** | GTACACAGGTCCCCATAGCA | mGM16441 |
| **SNP131.mMAGEB4.g10** | GTCCCCCTCCTGCTCTAATC | mMAGEB4 |
| **SNP131.mMAGEB4.g12** | GTTGCAAGCACCTCTACTGC | mMAGEB4 |
| **SNP131.mMAGEB4.g14** | AAAGTCTAAATCTCAAGGTG | mMAGEB4 |
| **SNP131.mMAGEB4.g15** | CCTTGAGATTTAGACTTTTG | mMAGEB4 |
| **SNP131.mMAGEB4.g16** | CTTGAGATTTAGACTTTTGA | mMAGEB4 |
| **SNP131.mMAGEB4.g18** | TCCAGAATCCTGATTAGAGC | mMAGEB4 |
| **SNP132.mMAGEB5.g11** | GAGACTCATCTTTTTGAGGA | mMAGEB5 |
| **SNP132.mMAGEB5.g13** | TGTGTTTCTTATTCTTCAAG | mMAGEB5 |
| **SNP132.mMAGEB5.g14** | TCTTATTCTTCAAGCGGTTG | mMAGEB5 |
| **SNP132.mMAGEB5.g15** | TATTCTTCAAGCGGTTGCGG | mMAGEB5 |
| **SNP132.mMAGEB5.g16** | GTGCTTATTCTTCTGACCCC | mMAGEB5 |
| **SNP133.mGM14781.g11** | GAGACTCATCTTTTTGAGGA | mGM14781 |
| **SNP133.mGM14781.g13** | TGTGTTTCTTATTCTTCAAG | mGM14781 |
| **SNP133.mGM14781.g14** | TCTTATTCTTCAAGCGGTTG | mGM14781 |

| | | |
|---|---|---|
| **SNP133.mGM14781.g15** | TATTCTTCAAGCGGTTGCGG | mGM14781 |
| **SNP133.mGM14781.g16** | GTGCTTATTCTTCTGACCCC | mGM14781 |
| **SNP134.mMAGEB6PS.g24** | TCTGGTCACACTATAGGGAC | mMAGEB6PS |
| **SNP134.mMAGEB6PS.g25** | CTGGTCACACTATAGGGACT | mMAGEB6PS |
| **SNP134.mMAGEB6PS.g27** | TTAAGTGGCCCCATTATTTC | mMAGEB6PS |
| **SNP134.mMAGEB6PS.g3** | GGACAGCCTAGACCCAATAC | mMAGEB6PS |
| **SNP134.mMAGEB6PS.g33** | AGGGTAAAGTGGGCCTGTAT | mMAGEB6PS |
| **SNP135.mMAGEB7PS.g1** | GCCCAGGGGACAGAAGAGTA | mMAGEB7PS |
| **SNP135.mMAGEB7PS.g17** | ACTTGCCGCAGCGTCTCCAG | mMAGEB7PS |
| **SNP135.mMAGEB7PS.g18** | CTTGCCGCAGCGTCTCCAGA | mMAGEB7PS |
| **SNP135.mMAGEB7PS.g3** | GGGACAGAAGAGTAAGGTCC | mMAGEB7PS |
| **SNP135.mMAGEB7PS.g4** | AACTAAGTTCCAGAACCTTG | mMAGEB7PS |
| **SNP135.mMAGEB7PS.g9** | TTTGCCCTCTGGAGACGCTG | mMAGEB7PS |
| **SNP136.mMAGEB8PS.g1** | TGCCTCGGGGTCAGAAAAAA | mMAGEB8PS |
| **SNP136.mMAGEB8PS.g13** | CTCCATCTTCTCCTCATTTG | mMAGEB8PS |
| **SNP136.mMAGEB8PS.g17** | TCATAGACTCAGCGGTAGAA | mMAGEB8PS |
| **SNP136.mMAGEB8PS.g23** | TGGGTAGTAACCCCCAAATG | mMAGEB8PS |
| **SNP136.mMAGEB8PS.g38** | AGCCTTTTTTTCTGACCCCG | mMAGEB8PS |
| **SNP137.mMAGEB10PS.g1** | TTGCCACAGCCGTCATGCCC | mMAGEB10PS |
| **SNP137.mMAGEB10PS.g11** | CAGGATGCTCAAGCCAAGGC | mMAGEB10PS |
| **SNP137.mMAGEB10PS.g18** | CCTGGGCCTCATCCTGGACC | mMAGEB10PS |
| **SNP137.mMAGEB10PS.g2** | TGCCACAGCCGTCATGCCCA | mMAGEB10PS |
| **SNP137.mMAGEB10PS.g23** | CTTTTGTCCCCTGGGCATGA | mMAGEB10PS |
| **SNP137.mMAGEB10PS.g24** | TCCCCTGGGCATGACGGCTG | mMAGEB10PS |
| **SNP137.mMAGEB10PS.g3** | GCCACAGCCGTCATGCCCAG | mMAGEB10PS |
| **SNP137.mMAGEB10PS.g4** | GCCCAGGGGACAAAAGAGTA | mMAGEB10PS |
| **SNP137.mMAGEB10PS.g6** | TGAGAAGCGTCGCCTGGTCC | mMAGEB10PS |
| **SNP137.mMAGEB10PS.g7** | GCGTCGCCTGGTCCAGGATG | mMAGEB10PS |
| **SNP137.mMAGEB10PS.g8** | CCTGGTCCAGGATGAGGCCC | mMAGEB10PS |
| **SNP138.mMAGEB16.g10** | ACATGGTTCCTCCACATCCT | mMAGEB16 |
| **SNP138.mMAGEB16.g21** | TGCTGCAACCTGTAGTTTCT | mMAGEB16 |
| **SNP138.mMAGEB16.g22** | TAGTTTCTGGGTCTCTTCAC | mMAGEB16 |

| | | |
|---|---|---|
| **SNP139.mLOC108168446.g15** | CCTCCTCCTCCACAGTGGGC | mLOC108168446 |
| **SNP139.mLOC108168446.g18** | GCTGGGCACCCTGGAGATTC | mLOC108168446 |
| **SNP139.mLOC108168446.g19** | TCCTCGAGTCTGCTGCCTTT | mLOC108168446 |
| **SNP139.mLOC108168446.g21** | GCTGCCTTTTGGCCCTGGAG | mLOC108168446 |
| **SNP139.mLOC108168446.g3** | ACAAGAGCAAGGGCCGCTCC | mLOC108168446 |
| **SNP139.mLOC108168446.g6** | GCCAAAAGGCAGCAGACTCG | mLOC108168446 |
| **SNP140.mGM5072.g19** | GGCCCCAGCTCCTCTGATAG | mGM5072 |
| **SNP140.mGM5072.g26** | AGTGCCTCTATCAGAGGAGC | mGM5072 |
| **SNP140.mGM5072.g30** | CCATCTCCCTGGTCAAGAGC | mGM5072 |
| **SNP140.mGM5072.g8** | GGAGAGAAGCCTAAACTCCA | mGM5072 |
| **SNP142.mMAGEB18.g21** | CTCCTGTGCATTTTTAGCAC | mMAGEB18 |
| **SNP142.mMAGEB18.g22** | GTGCATTTTTAGCACGGGCC | mMAGEB18 |
| **SNP142.mMAGEB18.g23** | CATTTTTAGCACGGGCCTGG | mMAGEB18 |
| **SNP142.mMAGEB18.g24** | GGGCCTGGCGGCGCTTCTCA | mMAGEB18 |
| **SNP142.mMAGEB18.g27** | GGGCACGGAGTTTGCTTTTC | mMAGEB18 |
| **SNP143.m4930550L24RIK.g17** | CTTTGCTTGATGCAGACGCT | m4930550L24RIK |
| **SNP143.m4930550L24RIK.g22** | CAGAGGATGAGGCGGTTTCC | m4930550L24RIK |
| **SNP143.m4930550L24RIK.g3** | AGGCTTTGAGGACCAGCTAG | m4930550L24RIK |
| **SNP143.m4930550L24RIK.g4** | CTTTGAGGACCAGCTAGAGG | m4930550L24RIK |
| **SNP144.mMAGED1.g18** | GGGACTAGCATTTGGCCCAC | mMAGED1 |
| **SNP144.mMAGED1.g2** | GGAAGCCATCCAGATCTCCG | mMAGED1 |
| **SNP144.mMAGED1.g26** | TGGGCGGAGCCTCGGAGATC | mMAGED1 |
| **SNP144.mMAGED1.g27** | CGGAGCCTCGGAGATCTGGA | mMAGED1 |
| **SNP144.mMAGED1.g28** | GATCTGGATGGCTTCCATCA | mMAGED1 |
| **SNP145.mMAGED2.g18** | TTTGAGACCTCTGTCGCCTT | mMAGED2 |
| **SNP145.mMAGED2.g20** | TGTCGCCTTTGGGTTCCCAG | mMAGED2 |
| **SNP145.mMAGED2.g24** | TTTCTGAGACCTCCAAGTTC | mMAGED2 |
| **SNP146.mMAGED3.g34** | GCCGGTGTGGTCTTATCCAT | mMAGED3 |
| **SNP146.mMAGED3.g36** | CTTGTTTTTCTTAGATCGGC | mMAGED3 |
| **SNP146.mMAGED3.g37** | TTCTTAGATCGGCTGGCAAT | mMAGED3 |
| **SNP146.mMAGED3.g41** | GTGGTCTCCGACTGTACATC | mMAGED3 |
| **SNP146.mMAGED3.g5** | GAAGAATTCTATTAAGCCTA | mMAGED3 |

| | | |
|---|---|---|
| **SNP147.mMAGEE1.g3** | CGCGCCGCCGCCGCGGTGGA | mMAGEE1 |
| **SNP147.mMAGEE1.g38** | CCCGTTGTTTCTGCGCGCAT | mMAGEE1 |
| **SNP147.mMAGEE1.g4** | GCGCCGCCGCCGCGGTGGAA | mMAGEE1 |
| **SNP147.mMAGEE1.g40** | CATTGGCCCTTCCACCGCGG | mMAGEE1 |
| **SNP147.mMAGEE1.g42** | CGCGGCGGCGGCGCGAATTC | mMAGEE1 |
| **SNP148.mMAGEE2.g1** | CGCCGATTACAGCAACAGCC | mMAGEE2 |
| **SNP148.mMAGEE2.g31** | CTGGCTGTTGCTGTAATCGG | mMAGEE2 |
| **SNP148.mMAGEE2.g33** | CGCTGCTTTGGCGCGCATTC | mMAGEE2 |
| **SNP148.mMAGEE2.g4** | ATGCAGGCTACTAATGCCTC | mMAGEE2 |
| **SNP148.mMAGEE2.g8** | TGAGATTCCAAACGATCCAC | mMAGEE2 |
| **SNP149.mMAGEF1.g22** | TAGGGAAGCCGCCAGGACCT | mMAGEF1 |
| **SNP149.mMAGEF1.g24** | GACCTGGGCTCGCCCTCGGC | mMAGEF1 |
| **SNP149.mMAGEF1.g25** | CTGGGCTCGCCCTCGGCAGG | mMAGEF1 |
| **SNP149.mMAGEF1.g26** | GGCTCGCCCTCGGCAGGAGG | mMAGEF1 |
| **SNP149.mMAGEF1.g4** | GTCCTGCCGCCTCCTGCCGA | mMAGEF1 |
| **SNP150.mNSMCE3.g2** | AGAAGCCGAGGGGCCGCGGT | mNSMCE3 |
| **SNP150.mNSMCE3.g46** | CCTGCGTGCTGGGCCGACCG | mNSMCE3 |
| **SNP150.mNSMCE3.g47** | CTGGGCCGACCGCGGCCCCT | mNSMCE3 |
| **SNP150.mNSMCE3.g5** | GCACGCAGGCTGACCCGGAG | mNSMCE3 |
| **SNP150.mNSMCE3.g6** | CACGCAGGCTGACCCGGAGC | mNSMCE3 |
| **SNP151.m1700020D05RIK.g1** | GTCACATGTTGCAAAAAACG | m1700020D05RIK |
| **SNP151.m1700020D05RIK.g14** | CTGACGAAAGCTTGGCGACG | m1700020D05RIK |
| **SNP151.m1700020D05RIK.g15** | AAGCTTGGCGACGGGGTGCC | m1700020D05RIK |
| **SNP151.m1700020D05RIK.g24** | ACACCCGGGCAGCAGTCGCT | m1700020D05RIK |
| **SNP151.m1700020D05RIK.g7** | AGAGAGATCGCGAGAGCGTC | m1700020D05RIK |
| **SNP152.mMAGEH1.g1** | TGCCTCGGGGACGGAAGAGT | mMAGEH1 |
| **SNP152.mMAGEH1.g2** | GACGGAAGAGTCGGCGCCGC | mMAGEH1 |
| **SNP152.mMAGEH1.g3** | TCGGCGCCGCCGGAACGCAA | mMAGEH1 |
| **SNP152.mMAGEH1.g31** | CAGCTGCCTTTGCGTTCCGG | mMAGEH1 |
| **SNP153.mMAGEL2.g1** | GTCGCAGCTAAGTACGAATC | mMAGEL2 |
| **SNP153.mMAGEL2.g2** | TCGCAGCTAAGTACGAATCT | mMAGEL2 |
| **SNP153.mMAGEL2.g3** | CGCAGCTAAGTACGAATCTG | mMAGEL2 |

| | | |
|---|---|---|
| **SNP153.mMAGEL2.g39** | TCAGAACCGTAGGGCGGCTA | mMAGEL2 |
| **SNP153.mMAGEL2.g4** | GCAGCTAAGTACGAATCTGG | mMAGEL2 |
| **SNP154.hNDN.g1** | CGACCCTAACTTTGCAGCCG | hNDN |
| **SNP154.hNDN.g29** | TGTCCTGCATCTCACAGTCG | hNDN |
| **SNP154.hNDN.g30** | CATCTCACAGTCGGGGACCT | hNDN |
| **SNP154.hNDN.g33** | CTGCAAAGTTAGGGTCGCTC | hNDN |
| **SNP154.hNDN.g5** | GACAGCGATGCCGTTCCGGT | hNDN |
| **SNP74.hMAGEA1.g1** | GAGTCTGCACTGCAAGCCTG | hMAGEA1 |
| **SNP74.hMAGEA1.g17** | GATCCTCCCCAGAGTCCTCA | hMAGEA1 |
| **SNP74.hMAGEA1.g18** | CCATCAACTTCACTCGACAG | hMAGEA1 |
| **SNP74.hMAGEA1.g19** | CCTCTGTCGAGTGAAGTTGA | hMAGEA1 |
| **SNP74.hMAGEA1.g20** | TCGAGTGAAGTTGATGGTAG | hMAGEA1 |
| **SNP74.hMAGEA1.g26** | CGGAGGCTCCCTGAGGACTC | hMAGEA1 |
| **SNP74.hMAGEA1.g30** | AGGATCTGTTGACCCAGCAG | hMAGEA1 |
| **SNP74.hMAGEA1.g31** | GGATCTGTTGACCCAGCAGT | hMAGEA1 |
| **SNP75.hMAGEA2.g13** | TACTCTAGTGGAAGTTACCC | hMAGEA2 |
| **SNP75.hMAGEA2.g14** | ACTCTAGTGGAAGTTACCCT | hMAGEA2 |
| **SNP75.hMAGEA2.g16** | TCTAGTGGAAGTTACCCTGG | hMAGEA2 |
| **SNP75.hMAGEA2.g21** | GTGGGGAGGACTCGGTGAGT | hMAGEA2 |
| **SNP75.hMAGEA2.g22** | GGACTCGGTGAGTCGGCAGC | hMAGEA2 |
| **SNP75.hMAGEA2.g4** | AGGCCTTGAGGCCCGAGGAG | hMAGEA2 |
| **SNP77.hMAGEA3.g1** | CAGCACTGCAAGCCTGAAGA | hMAGEA3 |
| **SNP77.hMAGEA3.g10** | CCTGGGCCTGGTGGGTGCGC | hMAGEA3 |
| **SNP77.hMAGEA3.g17** | TCTAGTTGAAGTCACCCTGG | hMAGEA3 |
| **SNP77.hMAGEA3.g18** | AGTTGAAGTCACCCTGGGGG | hMAGEA3 |
| **SNP77.hMAGEA3.g24** | GAGGCTCCCTGAGGACTCTG | hMAGEA3 |
| **SNP77.hMAGEA3.g25** | GCTCCCTGAGGACTCTGGGG | hMAGEA3 |
| **SNP77.hMAGEA3.g28** | GGATCTGGTGACTCGGCAGC | hMAGEA3 |
| **SNP77.hMAGEA3.g31** | GACTTCAACTAGAGTAGAAG | hMAGEA3 |
| **SNP77.hMAGEA3.g33** | AACTAGAGTAGAAGAGGAGG | hMAGEA3 |
| **SNP77.hMAGEA3.g36** | CCTGCGCACCCACCAGGCCC | hMAGEA3 |
| **SNP78.hMAGEA4.g1** | GAGTCAGCACTGCAAGCCTG | hMAGEA4 |

| | | |
|---|---|---|
| **SNP78.hMAGEA4.g10** | CCTGGGCCTGGTGGGTGCAC | hMAGEA4 |
| **SNP78.hMAGEA4.g18** | GTGCCTGCTGCTGAGTCAGC | hMAGEA4 |
| **SNP78.hMAGEA4.g2** | CAGCACTGCAAGCCTGAGGA | hMAGEA4 |
| **SNP78.hMAGEA4.g21** | GGACCTGCTGACTCAGCAGC | hMAGEA4 |
| **SNP78.hMAGEA4.g24** | GGCACTTCCTCCAGGGTGCC | hMAGEA4 |
| **SNP78.hMAGEA4.g25** | GCACTTCCTCCAGGGTGCCA | hMAGEA4 |
| **SNP78.hMAGEA4.g4** | AGGCGTTGAGGCCCAAGAAG | hMAGEA4 |
| **SNP79.hMAGEA5.g10** | TGTGCAGGCTGCCACTACTG | hMAGEA5 |
| **SNP79.hMAGEA5.g14** | TCCTCCTCTCCTCTGGTCCC | hMAGEA5 |
| **SNP79.hMAGEA5.g23** | AGGTCCTCTCAAGAGTCCTC | hMAGEA5 |
| **SNP79.hMAGEA5.g25** | TGAGGACTCTTGAGAGGACC | hMAGEA5 |
| **SNP79.hMAGEA5.g3** | AGGCCTTGACACCCAAGAAG | hMAGEA5 |
| **SNP79.hMAGEA5.g39** | CCAGGCCCAGGGCCTCTTCT | hMAGEA5 |
| **SNP79.hMAGEA5.g40** | CAGGCCCAGGGCCTCTTCTT | hMAGEA5 |
| **SNP79.hMAGEA5.g42** | TGGGTGTCAAGGCCTTCCTC | hMAGEA5 |
| **SNP79.hMAGEA5.g7** | AGAAGAGGCCCTGGGCCTGG | hMAGEA5 |
| **SNP79.hMAGEA5.g8** | GAAGAGGCCCTGGGCCTGGT | hMAGEA5 |
| **SNP80.hMAGEA6.g14** | TACTCTAGTTGAAGTCACCC | hMAGEA6 |
| **SNP80.hMAGEA6.g17** | TCTAGTTGAAGTCACCCTGG | hMAGEA6 |
| **SNP80.hMAGEA6.g18** | AGTTGAAGTCACCCTGGGGG | hMAGEA6 |
| **SNP80.hMAGEA6.g19** | GACTTCAACTAGAGTAGAAG | hMAGEA6 |
| **SNP80.hMAGEA6.g21** | AACTAGAGTAGAAGAGGAGG | hMAGEA6 |
| **SNP81.hMAGEA8.g1** | GCAGAAGAGTCAGCGCTACA | hMAGEA8 |
| **SNP81.hMAGEA8.g18** | TCAGGACTCTGGGGAGGACT | hMAGEA8 |
| **SNP81.hMAGEA8.g26** | ATCTGCACATCCATAAGCCC | hMAGEA8 |
| **SNP81.hMAGEA8.g27** | TAAGCCCTGGTGCCTCTCCT | hMAGEA8 |
| **SNP81.hMAGEA8.g3** | CAGCGCTACAAGGCTGAGGA | hMAGEA8 |
| **SNP81.hMAGEA8.g4** | CAAGGCTGAGGAAGGCCTTC | hMAGEA8 |
| **SNP81.hMAGEA8.g5** | GAGGAAGGCCTTCAGGCCCA | hMAGEA8 |
| **SNP81.hMAGEA8.g6** | AGGCCTTCAGGCCCAAGGAG | hMAGEA8 |
| **SNP82.hMAGEA9.g10** | ACAGGAACCCACAGGCGAGG | hMAGEA9 |
| **SNP82.hMAGEA9.g18** | AAGTCCTCCCCAGAGTCCTC | hMAGEA9 |

| | | |
|---|---|---|
| **SNP82.hMAGEA9.g19** | AGTCCTCCCCAGAGTCCTCA | hMAGEA9 |
| **SNP82.hMAGEA9.g20** | CCTCCCCAGAGTCCTCAGGG | hMAGEA9 |
| **SNP82.hMAGEA9.g22** | TAAAGTGTAGTAGACGGAAA | hMAGEA9 |
| **SNP82.hMAGEA9.g24** | GTAGTAGACGGAAATGGAGG | hMAGEA9 |
| **SNP82.hMAGEA9.g26** | AAGCGCCTCCCTGAGGACTC | hMAGEA9 |
| **SNP82.hMAGEA9.g35** | TGGGTTCCTGTGCACCCATC | hMAGEA9 |
| **SNP83.hMAGEA9B.g10** | ACAGGAACCCACAGGCGAGG | hMAGEA9B |
| **SNP83.hMAGEA9B.g22** | AAGCGCCTCCCTGAGGACTC | hMAGEA9B |
| **SNP83.hMAGEA9B.g31** | TGGGTTCCTGTGCACCCATC | hMAGEA9B |
| **SNP83.hMAGEA9B.g35** | TGGGCTTCAAGGTCTTCATC | hMAGEA9B |
| **SNP83.hMAGEA9B.g6** | GGAGAGGACTTGGGCCTGAT | hMAGEA9B |
| **SNP84.hMAGEA10.g1** | TCAATCCCAAAGTGAGACAC | hMAGEA10 |
| **SNP84.hMAGEA10.g26** | TGGGATTGAAGATCTTCTTC | hMAGEA10 |
| **SNP84.hMAGEA10.g27** | GGCATGCAGCGCTGACGCTT | hMAGEA10 |
| **SNP84.hMAGEA10.g28** | CGCTGACGCTTTGGAGCTCG | hMAGEA10 |
| **SNP84.hMAGEA10.g6** | CGAGGGTGCACAGGCTCCCC | hMAGEA10 |
| **SNP85.hMAGEA11.g1** | TCAACAGAAAGAGCCCCATA | hMAGEA11 |
| **SNP85.hMAGEA11.g2** | CATATGGTCCACAACTACAG | hMAGEA11 |
| **SNP85.hMAGEA11.g7** | ACTGTAGTTGTGGACCATAT | hMAGEA11 |
| **SNP85.hMAGEA11.g8** | CTGTAGTTGTGGACCATATG | hMAGEA11 |
| **SNP85.hMAGEA11.g9** | ATGGGGCTCTTTCTGTTGAC | hMAGEA11 |
| **SNP86.hMAGEA12.g1** | GAGTCAGCACTGCAAGCCTG | hMAGEA12 |
| **SNP86.hMAGEA12.g12** | TGCGCAGGCTCCTGCTACTG | hMAGEA12 |
| **SNP86.hMAGEA12.g14** | CTCCTCCTCCTCTACTCTAG | hMAGEA12 |
| **SNP86.hMAGEA12.g16** | TCTAGTGGAAGTCACCCTGC | hMAGEA12 |
| **SNP86.hMAGEA12.g24** | GCTCCCTGAGGACTGTGGGG | hMAGEA12 |
| **SNP86.hMAGEA12.g3** | CAAGCCTGAGGAAGGCCTTG | hMAGEA12 |
| **SNP86.hMAGEA12.g30** | GGTGACTTCCACTAGAGTAG | hMAGEA12 |
| **SNP86.hMAGEA12.g31** | GACTTCCACTAGAGTAGAGG | hMAGEA12 |
| **SNP86.hMAGEA12.g33** | CACTAGAGTAGAGGAGGAGG | hMAGEA12 |
| **SNP86.hMAGEA12.g41** | TGCTGACTCCTCTGCTCAAG | hMAGEA12 |
| **SNP87.hMAGEB1.g1** | TGCTCGTGAGAAACGCCGCA | hMAGEB1 |

| | | |
|---|---|---|
| **SNP87.hMAGEB1.g16** | TTCTGGGGAATGCCAGCAGC | hMAGEB1 |
| **SNP87.hMAGEB1.g18** | CTGGGGAATGCCAGCAGCAG | hMAGEB1 |
| **SNP87.hMAGEB1.g22** | AGTATCCCCTAAAACAGGAG | hMAGEB1 |
| **SNP87.hMAGEB1.g28** | CGTGAGCAACCTTGAGACCC | hMAGEB1 |
| **SNP87.hMAGEB1.g31** | TGCGGCGTTTCTCACGAGCA | hMAGEB1 |
| **SNP87.hMAGEB1.g4** | AAGGCGCGAGAGGAGACCCA | hMAGEB1 |
| **SNP87.hMAGEB1.g9** | CTCCTCCTCTCCTGTTTTAG | hMAGEB1 |
| **SNP88.hMAGEB2.g2** | GCAAGGCCCGAGATGAGACC | hMAGEB2 |
| **SNP88.hMAGEB2.g21** | TGAGACCCCGGGTCTCATCT | hMAGEB2 |
| **SNP88.hMAGEB2.g22** | GAGACCCCGGGTCTCATCTC | hMAGEB2 |
| **SNP88.hMAGEB2.g26** | TGCGGCGTTTCTCACGGGCA | hMAGEB2 |
| **SNP88.hMAGEB2.g4** | AAGGCCCGAGATGAGACCCG | hMAGEB2 |
| **SNP88.hMAGEB2.g5** | CCGGGGTCTCAATGTTCCTC | hMAGEB2 |
| **SNP89.hMAGEB3.g13** | GTGGTGGATAGTGCTCTCTG | hMAGEB3 |
| **SNP89.hMAGEB3.g15** | ATAGTAGCCCCCAAAATAAG | hMAGEB3 |
| **SNP89.hMAGEB3.g17** | AGTAGCCCCCAAAATAAGAG | hMAGEB3 |
| **SNP89.hMAGEB3.g22** | GGTGATCCTGGGTCTGACCC | hMAGEB3 |
| **SNP89.hMAGEB3.g25** | GCTGGCGTTTCTCACGTGCA | hMAGEB3 |
| **SNP89.hMAGEB3.g26** | AGCGTACTCTTCTGACCCCG | hMAGEB3 |
| **SNP89.hMAGEB3.g3** | GAGAAACGCCAGCAGACCCG | hMAGEB3 |
| **SNP89.hMAGEB3.g6** | GGTCAGACCCAGGATCACCA | hMAGEB3 |
| **SNP90.hMAGEB4.g10** | CCTGGGGAATGCCAAAAGCA | hMAGEB4 |
| **SNP90.hMAGEB4.g12** | AATGCCAAAAGCAAGGGAGC | hMAGEB4 |
| **SNP90.hMAGEB4.g18** | GCTGACCAACCTTGAGATCC | hMAGEB4 |
| **SNP90.hMAGEB4.g19** | CTGACCAACCTTGAGATCCT | hMAGEB4 |
| **SNP90.hMAGEB4.g25** | GCTGGCGTTTCTCACGGGCA | hMAGEB4 |
| **SNP90.hMAGEB4.g26** | AGCTTACTCTTCTGACCCCG | hMAGEB4 |
| **SNP90.hMAGEB4.g5** | CAGACCCAGGATCTCAAGGT | hMAGEB4 |
| **SNP91.hMAGEB5.g13** | GCTGCATGAACTCTCAGTGG | hMAGEB5 |
| **SNP91.hMAGEB5.g14** | CTGCATGAACTCTCAGTGGA | hMAGEB5 |
| **SNP91.hMAGEB5.g17** | GTGAGACCTCTGAGGAACAT | hMAGEB5 |
| **SNP91.hMAGEB5.g2** | TTAATGCAGGATCTGACGAA | hMAGEB5 |

| | | |
|---|---|---|
| **SNP91.hMAGEB5.g3** | TAATGCAGGATCTGACGAAA | hMAGEB5 |
| **SNP92.hMAGEB6.g10** | CTTGTCTGGGTGATTGTCGT | hMAGEB6 |
| **SNP92.hMAGEB6.g11** | ACCCAGACAAGCGCGAGAAG | hMAGEB6 |
| **SNP92.hMAGEB6.g20** | GAGACCCTGTGGCTGACCAT | hMAGEB6 |
| **SNP92.hMAGEB6.g24** | AGCTTACTCTTGTGACCCCG | hMAGEB6 |
| **SNP92.hMAGEB6.g4** | TGGTCAGCCACAGGGTCTCA | hMAGEB6 |
| **SNP92.hMAGEB6.g8** | ATCCTCTTCTCGCGCTTGTC | hMAGEB6 |
| **SNP93.hMAGEB10.g25** | GGCCTGACGGCGTTTTTCCC | hMAGEB10 |
| **SNP93.hMAGEB10.g27** | AGTTTACTCTTCTGACCTCG | hMAGEB10 |
| **SNP93.hMAGEB10.g3** | GAAGAGTAAACTCCGTGCCA | hMAGEB10 |
| **SNP93.hMAGEB10.g4** | TGCCAGGGAAAAACGCCGTC | hMAGEB10 |
| **SNP93.hMAGEB10.g5** | GAAAAACGCCGTCAGGCCCG | hMAGEB10 |
| **SNP93.hMAGEB10.g6** | AAACGCCGTCAGGCCCGAGG | hMAGEB10 |
| **SNP94.hMAGEB16.g11** | GGAGCTTCCTTCAGTTTGCC | hMAGEB16 |
| **SNP94.hMAGEB16.g14** | TTTGCCAGGCACTAGAGGAT | hMAGEB16 |
| **SNP94.hMAGEB16.g21** | TGGAGACCTGTGCAACCTCC | hMAGEB16 |
| **SNP94.hMAGEB16.g25** | GGGTCTCACTGAAGGTCTGA | hMAGEB16 |
| **SNP94.hMAGEB16.g26** | TGCTGATCATGTGTGCATCG | hMAGEB16 |
| **SNP95.hMAGEB17.g1** | TGCCCGTGAGAAACGCCGCC | hMAGEB17 |
| **SNP95.hMAGEB17.g2** | GAGAAACGCCGCCAGGCTCG | hMAGEB17 |
| **SNP95.hMAGEB17.g28** | CTTGAGCACCCCCGAGACAT | hMAGEB17 |
| **SNP95.hMAGEB17.g29** | ATTGGTCTTCACCTCGAGCC | hMAGEB17 |
| **SNP95.hMAGEB17.g31** | GAGCCTGGCGGCGTTTCTCA | hMAGEB17 |
| **SNP95.hMAGEB17.g32** | AGCCTGGCGGCGTTTCTCAC | hMAGEB17 |
| **SNP95.hMAGEB17.g33** | GGCGGCGTTTCTCACGGGCA | hMAGEB17 |
| **SNP95.hMAGEB17.g34** | CGCTTACTCGCCTGACCGCG | hMAGEB17 |
| **SNP95.hMAGEB17.g5** | AGGTGAAGACCAATGTCTCG | hMAGEB17 |
| **SNP95.hMAGEB17.g7** | TCAAGCCACCGCAGCAGAGA | hMAGEB17 |
| **SNP96.hMAGEB18.g1** | TGCCCGTGAGAAACGCCACC | hMAGEB18 |
| **SNP96.hMAGEB18.g11** | TGACTCTCCTTCTGCCACAG | hMAGEB18 |
| **SNP96.hMAGEB18.g15** | AGCCTGGTGGCGTTTCTCAC | hMAGEB18 |
| **SNP96.hMAGEB18.g16** | GGTGGCGTTTCTCACGGGCA | hMAGEB18 |

| | | |
|---|---|---|
| **SNP96.hMAGEB18.g17** | AGCTTACTCTTCTGACCTCG | hMAGEB18 |
| **SNP96.hMAGEB18.g3** | TCGTTGTGAGAATCAGGATC | hMAGEB18 |
| **SNP96.hMAGEB18.g6** | GGGAGCTACGCAGGCCACTG | hMAGEB18 |
| **SNP97.hMAGEC1.g1** | GACAAGGATATGCCTACTGC | hMAGEC1 |
| **SNP97.hMAGEC1.g12** | ACAACTCTGAGGACTCTCAG | hMAGEC1 |
| **SNP97.hMAGEC1.g14** | GAGGAACTCTGGAGAAGACT | hMAGEC1 |
| **SNP97.hMAGEC1.g2** | ACAAGGATATGCCTACTGCT | hMAGEC1 |
| **SNP98.hMAGEC2.g18** | TCGTTGTCAACGTTGCGGAA | hMAGEC2 |
| **SNP98.hMAGEC2.g19** | ACGTTGCGGAATGGAACGCC | hMAGEC2 |
| **SNP98.hMAGEC2.g20** | GCGGAATGGAACGCCTGGAA | hMAGEC2 |
| **SNP98.hMAGEC2.g21** | CGGAATGGAACGCCTGGAAC | hMAGEC2 |
| **SNP98.hMAGEC2.g22** | AATGGAACGCCTGGAACGGG | hMAGEC2 |
| **SNP99.hMAGEC3.g1** | GATGAGGATATGCCTGCTGC | hMAGEC3 |
| **SNP99.hMAGEC3.g11** | GCAGGAGTCTAGAGGACTCT | hMAGEC3 |
| **SNP99.hMAGEC3.g13** | GAGTCTAGAGGACTCTGGGG | hMAGEC3 |
| **SNP99.hMAGEC3.g21** | CCCTGGGGAGAGATCTTGGG | hMAGEC3 |
| **SNP99.hMAGEC3.g8** | CCACAAAAGAGGGGATGAGC | hMAGEC3 |

| siRNA | Sequence (5' - 3') | |
|---|---|---|
| RevL1 | ACUACAUCGUGAUUCAAACUU | |
| PanMAGEA2 | GGUAAAGAUCAGUGGAGGA | |
| PanMAGEA4 | GGUCACAAAGGCAGAAAUG | |
| MAGEA3/6 | GAUGGUUGAAUGAGCGUCA | |
| MAGEA11 #1 | GGGAAAUGGGCCAAUGCAU | |
| MAGEA11 #4 | CUGAUAGACCCUGAGUCCU | |
| MAGEA12 #3 | CACCAUUUGCUAAAGAUCA | MAGEA12 (pool) |
| MAGEA12 #4 | CUAUACUCUCUGGAGUCAA | |
| MAGEA12 #5 | CAACUAUACUCUCUGGAGU | |
| MAGEA12 #6 | GUAUUGUUAGUAGUGAGUU | |
| MAGEB1 #1 | GUCAGUAAGCUAAACCUCA | |
| MAGEB1 #2 | CUUCAUUCUACGUAAGUAU | |
| MAGEB2 #7 | CCGUUACAAAGGGAGAAAU | |
| MAGEB2 #10 | CGUUACAAAGGGAGAAAUG | |
| MAGEB2 UTR #1 | GGCAGAUUCUUUACUUUGU | |
| MAGEB2 UTR #2 | GUGGUCAAUUCUUGGUUUA | |
| MAGEC1 #3 | CUCUUCUCCAGAUUCCUAU | |
| MAGEC1 #4 | CAGAGUUUCCCUGAGAGUU | |
| MAGEC2 #5 | GUGAAAGCCUCAGUGUCAU | |
| MAGEC2 #7 | GCAGUUUAGGUUCUAGGUA | |
